# Chromosome-level genomics and cross-species enzyme screening enable high-titer bakuchiol production in *Yarrowia lipolytica*

**DOI:** 10.64898/2026.11.11.765817

**Authors:** Jing Li, Hong Chang, Chuan Liu, Shuai Li, Jin Zhang, Jie Bai, Han Wang, Xiaonan Liu, Zhijin Fan, Huifeng Jiang

**Author notes:** Correspondence: Huifeng Jiang; Xiaonan Liu; Zhijin Fan.

## Abstract

The diversity of enzymes supporting bakuchiol biosynthesis remains incompletely characterized, limiting the repertoire available for microbial pathway engineering. Here, we combined chromosome-level genomics, cross-species enzyme screening and metabolic engineering to establish high-titer bakuchiol production in *Yarrowia lipolytica*. We assembled a 471.76-Mb *Psoralea corylifolia* genome anchored to 11 pseudochromosomes, with a contig N50 of 40.33 Mb. Genome-guided screening and functional validation identified four prenyltransferases from *P. corylifolia* and *Vigna radiata* whose N-terminally truncated variants supported bakuchiol production in yeast. These results extend experimental evidence for bakuchiol-producing activity across legume prenyltransferase sequences and species. For PcPT6, corresponding to the previously reported PcPT07, removing 40 N-terminal amino acids increased bakuchiol production by 53% relative to the full-length enzyme. We reconstructed bakuchiol biosynthesis in *Y. lipolytica* and reinforced aromatic precursor synthesis to increase *p*-coumaric acid supply. Coordinating mevalonate pathway reinforcement with increased *t40PcPT6* gene dosage further improved bakuchiol production. Following fermentation optimization, titers reached 108.29 mg/L in shake flasks and 6.03 g/L in a 7.5-L fed-batch bioreactor. These findings connect an expanded set of functional prenyltransferases with coordinated pathway engineering, enabling gram-per-liter microbial bakuchiol production.

## Introduction

*Psoralea corylifolia* L. (Fabaceae) is a medicinal plant whose fruits contain coumarins, flavonoids and monoterpene phenols (Chen et al., 2023). Bakuchiol (BAK), a characteristic seed-derived meroterpenoid, has attracted interest for its retinol-like properties and potential cosmetic and pharmaceutical applications. Reported activities include anti-aging, skin-whitening and anti-inflammatory effects, together with low skin irritation (Jafernik et al., 2021).

Plant extraction and chemical synthesis present distinct challenges for BAK supply. Extraction requires separation from co-occurring plant constituents, whereas chemical synthesis involves lengthy routes and challenges in stereoselective synthesis (Adarsh Krishna et al., 2022; Chen and Li, 2008). These constraints motivate microbial production routes that combine renewable feedstocks with genetically tractable hosts.

BAK biosynthesis connects phenylpropanoid and isoprenoid metabolism through the coupling of *p*-coumaric acid (*p*-CA) with geranyl diphosphate (GPP). Early isotope-labeling studies traced these precursor contributions to the aromatic moiety and monoterpene side chain, respectively (Banerji and Chintalwar, 1983; 1984). The prenyltransferase (PT) mediating this coupling is therefore central to pathway reconstruction. In a microbial host, its performance must be considered alongside the supply of both precursors.

Systematic identification of BAK-producing enzymes requires access to candidate genes beyond those readily detected in individual transcriptomes. Plant membrane-bound PTs can be difficult to characterize because of their membrane-associated architecture (Huang et al., 2014). Transcriptome mining previously identified PcPT07, which supported de novo BAK production in *Saccharomyces cerevisiae* (Zheng et al., 2024). Metabolic and enzyme engineering increased the reported titer to 9.28 mg/L. This work established the terminal biosynthetic step, while leaving the functional diversity of related PTs and their suitability for alternative hosts open to investigation. A chromosome-level genome can broaden the candidate pool and support comparisons across related species.

The oleaginous yeast *Yarrowia lipolytica* offers an alternative host for BAK biosynthesis, supported by established genetic resources and metabolic engineering tools (Chen et al., 1997). Its acetyl-CoA metabolism has been exploited for the production of flavonoids and terpenoids (Che et al., 2026; Gu et al., 2026). Its capacity to accommodate lipophilic compounds provides a further rationale for evaluating BAK production. Effective pathway reconstruction nevertheless requires coordination of aromatic precursor synthesis, GPP supply through the mevalonate (MVA) pathway and terminal PT-mediated conversion.

Here, we assembled a chromosome-level *P. corylifolia* genome and used it to guide PT screening in *P. corylifolia* and *Vigna radiata*. We evaluated candidate enzymes and their N-terminal truncations in yeast, then reconstructed BAK biosynthesis in *Y. lipolytica*. Iterative pathway engineering and fermentation optimization tested how precursor supply and terminal conversion could be coordinated to support high-titer BAK production.

## Results

### 1. Chromosome-scale genome assembly and annotation of *P. corylifolia*

Analysis of the k-mer (k = 21) frequency distribution estimated the *P. corylifolia* genome size at approximately 484.06 Mb, with 0.03% heterozygosity and 51.03% repeat content (Fig. S1). To construct a chromosome-level reference genome, we generated 39.3 Gb of PacBio HiFi long-read data. Following filtering and polishing, assembly with Hifiasm yielded a 471.76-Mb genome comprising 69 contigs, with a contig N50 of 40.33 Mb and a GC content of 35.90%, consistent with the genome survey (Fig. S2; Supplementary Table 1). We then used 100.86 Gb of high-quality Hi-C data to anchor the assembly to 11 pseudochromosomes (2*n* = 22) (Fig. 1B, Fig. S3). The final assembly comprised 63 scaffolds, with a scaffold N50 of 42.23 Mb (Supplementary Table 1). Assessment using Benchmarking Universal Single-Copy Orthologs (BUSCO v5.2.2) and the embryophyte dataset from OrthoDB 10 identified 98.33% of BUSCO genes as complete (1587 of 1614 genes; Supplementary Table 2). These metrics indicate high assembly contiguity and completeness.

**Fig. 1.**
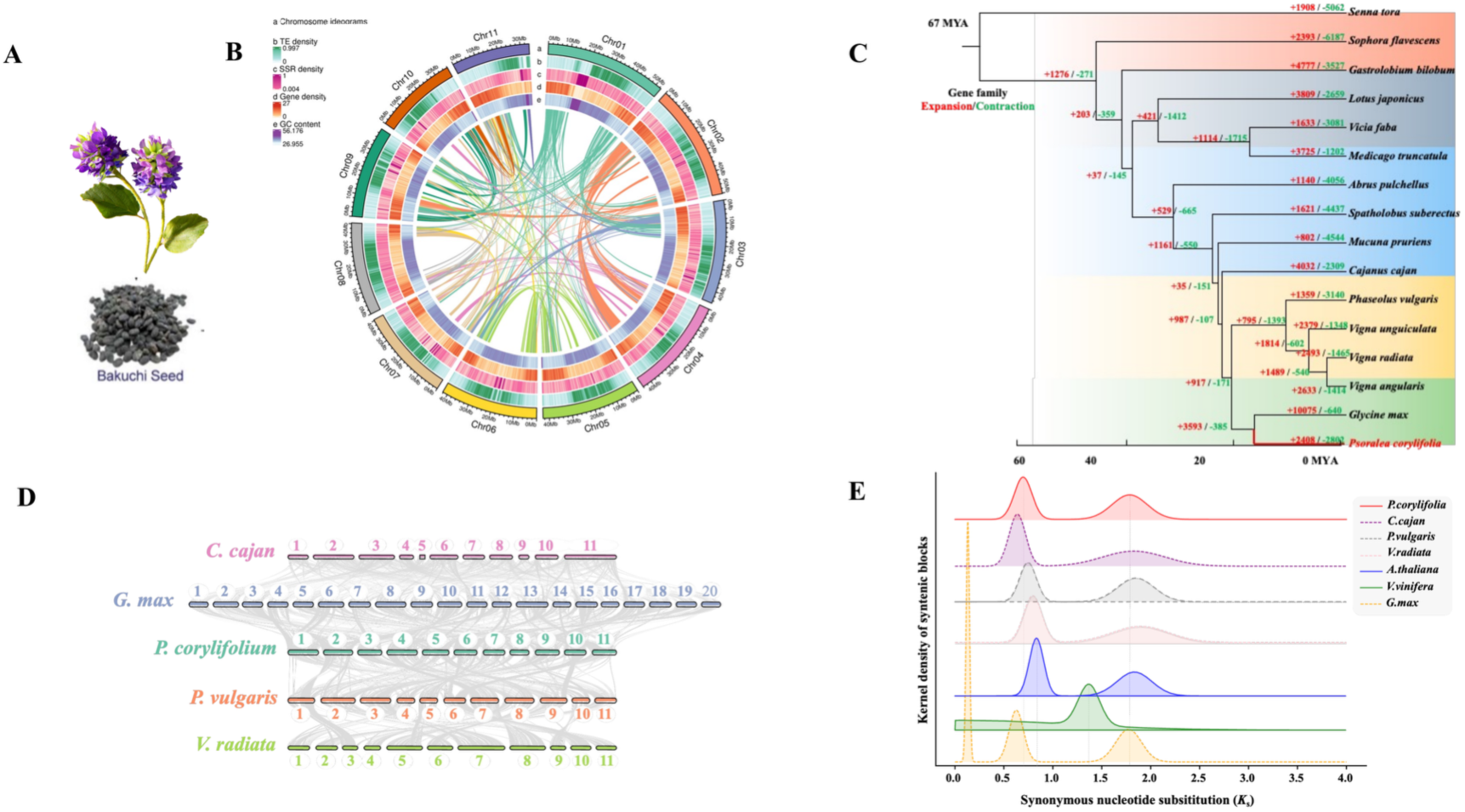
Analysis of genomic evolution of *Psoralea corylifolia*. (A) Morphological overview of *P. corylifolia* and its seeds. (B) Circos plot of the 11 *P. corylifolia* pseudochromosomes. From outside to inside, the tracks show chromosome ideograms, transposable element (TE) density, simple sequence repeat (SSR) density, gene density, and GC content. Central lines connect syntenic regions between chromosomes. Density tracks were calculated in 100-kb windows with a 10-kb step. (C) Phylogenetic tree, divergence times, and gene-family expansion/contraction for *P. corylifolia* and 15 related legume species. The tree was reconstructed from 117 single-copy orthologous genes using OrthoFinder2/STAG and calibrated with TimeTree. Numbers on branches indicate gene-family expansions/contractions, and the time scale is shown in million years ago (MYA). The topology places *Glycine max* as the closest relative of *P. corylifolia*. (D) Genome-wide collinear blocks among *P. corylifolia*, *Cajanus cajan*, *Phaseolus vulgaris*, *Vigna radiata*, and *Glycine max*. *P. corylifolia* shows one-to-one syntenic relationships with the three diploid legumes and a one-to-two relationship with the paleopolyploid *G. max*. (E) Distribution of synonymous substitution rates (Ks) for paralogous genes. The peaks at Ks ≈ 0.7 and ≈ 1.8 correspond to the legume-specific whole-genome duplication (LWGD) and the ancient gamma whole-genome triplication (gamma-WGT), respectively. *G. max* has an additional recent WGD peak at Ks ≈ 0.13.

We next annotated the repetitive elements and protein-coding genes. Approximately 49.60% (233.98 Mb) of the *P. corylifolia* genome was classified as transposable elements (TEs), among which long terminal repeat (LTR) retrotransposons were predominant, accounting for 89.77% of the total TEs (Supplementary Table 3). Among the 150,223 identified LTR elements, 72,882 (representing 30.66% of the genome) belonged to the Copia superfamily, while 21,469 (6.23% of the genome) were assigned to the Gypsy superfamily (Supplementary Table 3). Integrating *ab initio* prediction, homology-based searches, and transcriptomic evidence from leaves, stems, seeds, and roots identified 31,907 protein-coding genes (Supplementary Table 4). The mean gene, coding sequence, and intron lengths were 3713.69, 1439.85, and 1980.19 bp, respectively (Supplementary Table 5). Of the predicted genes, 99.18% were located on the 11 pseudochromosomes.

To characterize the predicted gene set, we compared the protein-coding genes with those of *Arabidopsis thaliana* and six Fabaceae species, including *Mucuna pruriens, Phaseolus vulgaris, Glycine max, Spatholobus suberectus, Vigna angularis, and Vigna radiata*, and identified 21,299 orthologous groups covering 497,631 genes in these eight species. Functional annotation revealed that 99.02% of the predicted *P. corylifolia* genes matched entries in at least one public database, including NR, eggNOG, Gene Ontology (GO), Kyoto Encyclopedia of Genes and Genomes (KEGG), TrEMBL, KOG, Swiss-Prot, and Pfam (Supplementary Table 6). Annotation of non-coding RNAs (ncRNAs) identified 499 transfer RNAs (tRNAs), 15,814 ribosomal RNAs (rRNAs), 80 small nuclear RNAs (snRNAs), 336 small nucleolar RNAs (snoRNAs) and 102 microRNAs (miRNAs) (Supplementary Table 7). Together, the assembly and annotation provide a genomic resource for investigating BAK biosynthesis.

### 2. Phylogenomic and evolutionary analysis of *P. corylifolia*

To resolve the evolutionary position of *P. corylifolia* within the Fabaceae, orthologous protein sequences from *P. corylifolia* and 15 other species with published genomes (*Abrus pulchellus, Cajanus cajan, Gastrolobium bilobum, G. max, Lotus japonicus, Medicago truncatula, M. pruriens, P. vulgaris, Senna tora, Sophora flavescens, S. suberectus, Vicia faba, V. angularis, V. radiata and Vigna unguiculata*) were used to reconstruct phylogenetic relationships and estimate divergence times. A total of 28,582 orthogroups were identified across these 16 genomes, including 117 single-copy orthologous genes and a highly conserved core set of 11,316 (Fig.S5) present in all species. Using the multi-copy-aware STAG algorithm implemented in OrthoFinder2, we reconstructed their species tree (Figs. S4; Supplementary Tables 8 and 9). The resulting topology placed soybean (*G. max*) as the sister lineage to *P. corylifolia*, with *Vigna* species more distantly related (Fig. 1C). Analysis the previous species tree of single-copy orthologs with time calibrations from TimeTree estimated that *P. corylifolia* diverged from *G. max* approximately 14.1 million years ago (MYA) and from *Vigna* species approximately 23.82 MYA, with *S. tora* serving as the outgroup (Fig. 1C).

To examine chromosomal conservation and rearrangements within the Fabaceae, we performed genome-wide synteny analysis. The 11 *P. corylifolia* pseudochromosomes were compared with four representative legume genomes with available chromosome-level assemblies: *C. cajan* (*n* = 11), *P. vulgaris* (*n* = 11), *V. radiata* (*n* = 11), and the palaeopolyploid *G. max* (*n* = 20). Collinearity analyses revealed a one-to-one (1:1) relationship between *P. corylifolia* and the three diploid legumes (*C. cajan, P. vulgaris,* and *V. radiata*). Large contiguous collinear blocks covered most of their respective chromosome arms, while collinearity with *P. vulgaris* and *V. radiata* was interrupted in several regions (Fig. 1D). This 1:1 correspondence is consistent with broad conservation of the ancestral legume karyotype, accompanied by chromosomal rearrangements since divergence from the common ancestor. In comparison with its sister lineage *G. max* (*n* = 20), *P. corylifolia* displayed a one-to-two (1:2) syntenic mapping pattern. Most chromosomal regions in *P. corylifolia* could be mapped to two duplicated homologous regions across the soybean chromosomes (Fig. 1D). This pattern is consistent with a soybean-specific whole-genome duplication (WGD) followed by chromosomal rearrangements.

We next analyzed synonymous substitutions per synonymous site (*Ks*) for paralogous genes in collinear regions of the five legume species, *A. thaliana* and *Vitis vinifera*. For *P. corylifolia* and the other diploid legumes (*P. vulgaris, C. cajan,* and *V. radiata*), the *Ks* distributions consistently showed two peaks: a primary peak at *Ks* ≈ 0.7 corresponding to the core legume-specific whole-genome duplication (LWGD) event inherited from their common ancestor, while *A. thaliana* also displayed a similar peak representing the Brassicaceae-specific alpha and beta duplication events (α/β-WGD). A secondary peak also appeared at Ks ≈ 1.8 in all four species; these peaks are consistent with the ancient core-eudicot gamma whole-genome triplication (γ-WGT) event (Fig. 1E). Concurrently, *V. vinifera* showed a single peak at Ks ≈ 1.4, consistent with the ancestral γ-WGT event. In contrast, the paleopolyploid *G. max* displayed three peaks, including a lineage-specific recent WGD peak at *Ks* ≈ 0.13 in addition to the LWGD peak and γ-WGT peak (Fig. 1E). Together, these *Ks* profiles support retention of the ancestral LWGD signature in *P. corylifolia*, without evidence of the additional recent WGD observed in soybeans. These analyses establish the comparative genomic context for investigating the evolution of *P. corylifolia* prenyltransferases.

### 3. Genome mining and functional characterization of PcPT6 and related legume prenyltransferases

Bakuchiol (BAK) primarily accumulates in the seeds of *P. corylifolia* (Nizam et al., 2023). Identifying the prenyltransferases that modify the phenolic precursor is central to elucidating BAK biosynthesis. Previous studies (Zheng *et al*., 2024) have identified prenyltransferases from *P. corylifolia* via transcriptome-based analysis, enabling *de novo* BAK synthesis from glucose in *S. cerevisiae*. However, RNA-Seq-based screening may overlook genes with low or condition-dependent expression. To address this limitation, we integrated genome-wide multi-omics screening with functional validation to systematically assessment and identify additional *P. corylifolia* prenyltransferases.

An HMMER search targeting the UbiA-type PT domain (PF01040) was performed against the high-quality *P. corylifolia* genome, identifying 32 putative PT sequences. Of these, 10 sequences, likely representing transcript variants, pseudogenes, or truncated fragments, were excluded, leaving 22 high-confidence candidates. The remaining candidates were included in a phylogenetic analysis with 67 previously characterized plant PTs, with their tissue-specific expression profiles (FPKM) mapped onto the phylogenetic tree (Fig. S6, Supplementary Table 10). Phylogenetic analysis revealed that the 22 PTs were distributed across six evolutionary distinct clades. The isoflavonoid and stilbenoid clades contained the most *P. corylifolia* PTs, suggesting expansion of these groups. Characterized enzymes in the isoflavonoid-associated group modify genistein, chalcone, and flavone scaffolds, whereas those in the stilbenoid group utilize resveratrol. We also prioritized candidates associated with the tocopherol, tocotrienol, xanthone, and shikonin groups because their substrates share structural features with *p*-CA. Expression profiles narrowed the selection to 10 genes highly expressed in seeds, the primary site of BAK accumulation. We then used AlphaFold3 structure predictions to assess substrate-binding pockets and putative catalytic residues, selecting six candidates for experimental validation.

These six candidate genes were codon-optimized for expression in *S. cerevisiae* and individually cloned into the YCplac22 vector. To increase the availability of the prenyl donor geranyl diphosphate (GPP), the F96W/N127W double mutant of the endogenous *S. cerevisiae ERG20* gene (*ScERG20*^WW^) (Ignea et al., 2014), which was previously reported to enhance GPP production, was generated and cloned into the YCplac33 vector. The resulting plasmids carrying *ScERG20*^WW^ and each candidate prenyltransferase were co-transformed into *S. cerevisiae* W303-1B to generate strains ScP01-ScP06, whereas strain ScP00 harboring the corresponding empty vectors served as a negative control. Following cultivation with 1.5 mM *p*-CA, HPLC analysis revealed that strain ScP06, expressing *PcPT6*, produced a new metabolite with a retention time of 22.9 min, matching that of an authentic BAK standard (Fig. 2A). Mass spectrometric analysis further confirmed that the molecular mass of this product matched that of BAK (Fig. S7B), supporting the role of PcPT6 in converting *p*-CA to BAK.

**Fig. 2.**
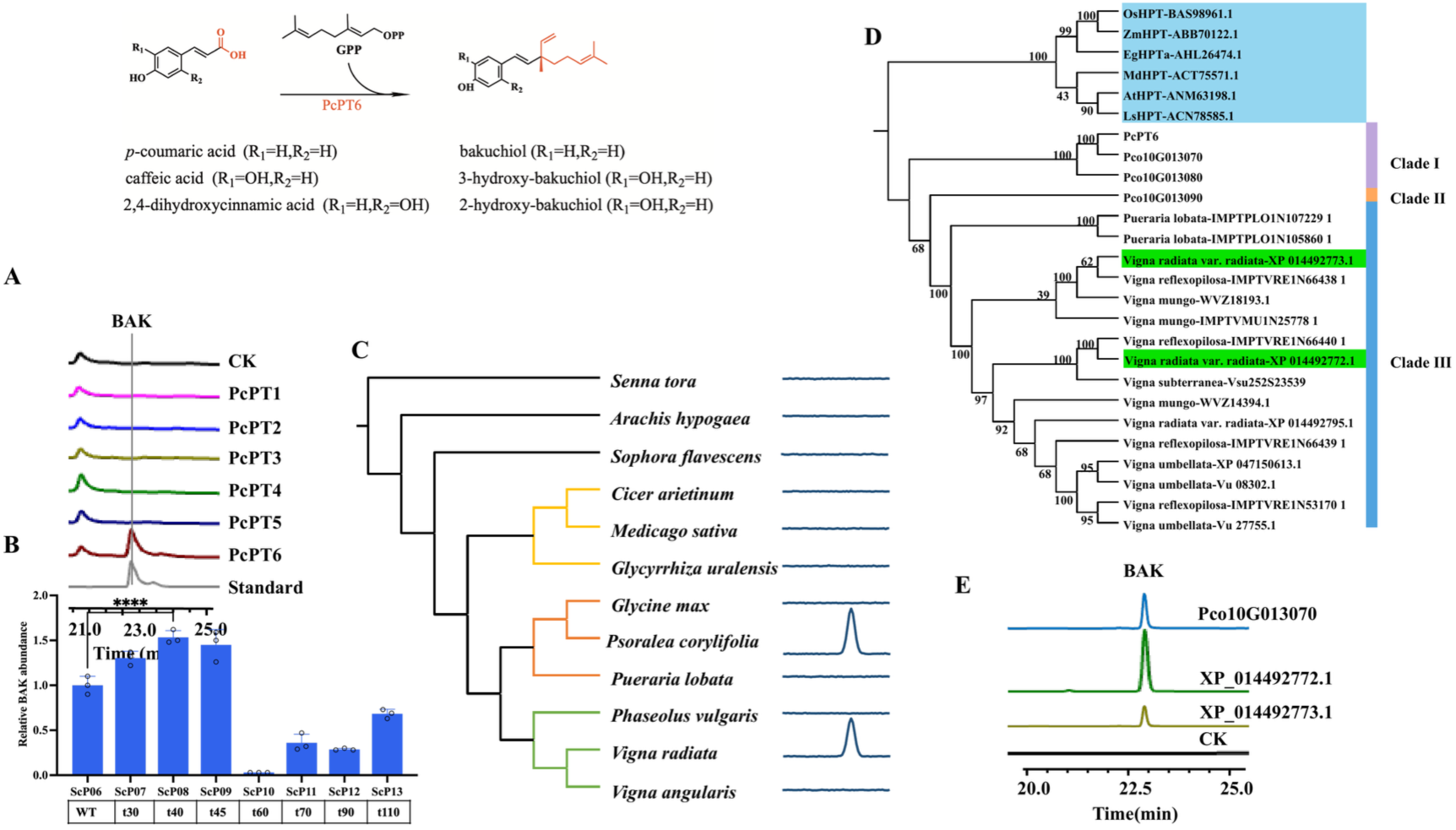
Functional identification of an UbiA-type prenyltransferase involved in BAK biosynthesis. (A) HPLC chromatograms of *Saccharomyces cerevisiae* strains ScP00 (control) and ScP01-ScP06 expressing PcPT1-PcPT6, together with an authentic BAK standard. Strain ScP06 produced a product with a retention time matching BAK (22.9 min). (B) Relative BAK abundance in strains expressing full-length PcPT6 or N-terminal truncation variants t30-t110. Truncation of 40 amino acids (t40PcPT6) gave the highest BAK production, which was 53% higher than that of the full-length enzyme. (C) Analysis of BAK in the seeds of representative Fabaceae species. BAK was detected in *Vigna radiata*, whereas no corresponding peak was detected in *Glycine max* or *Vigna angularis*. (D) Phylogenetic tree of PcPT6 homologs from representative Fabaceae species. Tocopherol HPTs were used as the outgroup, and the PcPT6 clade is divided into Clades I-III; Clades I and II contain *P. corylifolia* sequences, whereas Clade III contains sequences from *Pueraria lobata* and *Vigna* species. (E) HPLC analysis of BAK production. Truncated Cco10G013070 and the two of the *V. radiata* PTs (XP_014492772.1 and XP_014492773.1) produced detectable BAK; CK, control. Data in (B) represent the mean of n = 3 biologically independent samples, and error bars show standard deviation (s.d.). Circles represent individual raw data points. Statistical analysis in (B) was carried out using Student’s t-test with Welch’s correction (one-tailed; two-sample unequal variance; *P < 0.05, **P < 0.01, ***P < 0.001, ****P < 0.0001), for the comparison indicated by the bracket.

We next investigated the substrate selectivity of PcPT6 using a panel of structural analogs of *p*-CA. In addition to *p*-CA, PcPT6 converted caffeic acid into a product whose mass spectrum matched that of an authentic 3-hydroxybakuchiol standard (Fig. S7C). The new product obtained after feeding 2,4-dihydroxycinnamic acid was putatively assigned as 2-hydroxybakuchiol based on its mass spectrum, the substrate structure, and the catalytic characteristics of PcPT6 (Fig. S7D). This structural assignment remains tentative because an authentic 2-hydroxybakuchiol standard was unavailable. In contrast, no detectable prenylated products were observed when other tested substrates, including ferulic acid, were supplied. These results indicate that PcPT6 exhibits a moderate substrate promiscuity toward hydroxycinnamic acids and favors the tested phenylpropanoid substrates containing both a para-hydroxyl group and a carboxyl group. The lack of detectable activity toward ferulic acid suggests that its additional 3-*O*-methyl group may interfere with substrate recognition or productive positioning within the active site. These results define the substrate range of PcPT6 within the tested panel.

Plant UbiA-family prenyltransferases are generally integral membrane proteins associated with plastid membranes and often contain N-terminal plastid transit peptides that direct their targeting to chloroplasts (Munakata and Yazaki, 2024). These peptides are subsequently cleaved during plastid import, whereas heterologous hosts such as *S. cerevisiae* lack both plastids and the corresponding processing machinery. N-terminal truncation has therefore been widely used to improve the heterologous expression and activity of plant membrane-associated prenyltransferases (Guo et al., 2023; Yang et al., 2024). To determine whether this strategy could enhance PcPT6 activity, a series of truncation mutants lacking 30-110 amino acids from the N terminus were constructed and individually expressed in *S. cerevisiae* W303-1B. The resulting strains were cultivated with 1.5 mM *p*-CA, and PcPT6 activity was evaluated based on BAK production. Truncations of 30-45 amino acids substantially enhanced BAK production, with the 40-amino-acid truncation mutant, designated t40PcPT6, yielding the highest BAK titer and producing 53% more BAK than the full-length enzyme (Fig. 2B). In contrast, further truncation sharply reduced BAK production, which was nearly abolished when 60 amino acids were removed. These results identify a narrow N-terminal truncation window that improves PcPT6 performance in yeast. The loss of BAK production after more extensive truncation may reflect impaired protein stability, membrane insertion, or catalytic function.

To determine whether other legumes can produce BAK, we analyzed seeds from 11 representative Fabaceae species. BAK was detected in *V. radiata* seeds, whereas no corresponding peak was detected in *G. max* or *V. angularis* (Fig. 2C). We then identified homologous prenyltransferases from the genomes of these 11 species, together with four additional *Vigna* species, and constructed a phylogenetic tree using tocopherol HPTs as the outgroup. The resulting tree resolved the PcPT6 clade into three subclades: Clades I and II comprised PTs exclusively from *P. corylifolia*, whereas Clade III included PTs from *Pueraria lobata*, *V. radiata*, *V. mungo*, *V. reflexopilosa*, *V. umbellata*, and *V. subterranea* (Fig. 2D). We selected three representative PTs from *V. radiata* in Clade III, along with three additional PTs from Clades I and II for functional characterization. Following N-terminal truncation analogous to that used for PcPT6, truncated Cco10G013070 and the two of the three *V. radiata* PTs (XP_014492772.1 and XP_014492773.1) supported detectable BAK production (Fig. 2E). Comparison based on BAK production showed that t40PcPT6 outperformed *V. radiata* enzymes. Accordingly, t40PcPT6 was selected for subsequent metabolic engineering and BAK production experiments.

Together, genome-wide mining and functional screening identified PcPT6 as a BAK-producing prenyltransferase and revealed related functional enzymes in *V. radiata*. The 53% increase in BAK production achieved with t40PcPT6 supports its use in microbial pathway engineering.

### 4. Stepwise engineering of the *p*-CA pathway

To reconstruct the BAK biosynthetic pathway in *Y. lipolytica*, we introduced precursor biosynthesis modules for *p*-CA and GPP and systematically evaluated their contributions to BAK production. Strain YL00, previously constructed in our laboratory, was used as the parental strain, and a CRISPR-Cas9-based marker-free genome integration strategy was employed for pathway assembly (Wang et al., 2022). Because *p*-CA can be synthesized through either the tyrosine-or phenylalanine-derived route, we simultaneously reinforced both pathways by introducing five genes, *CluTAL, RgPAL, EbC4H, ATR2,* and *CYB5*. At this stage, full-length *PcPT6*, codon-optimized for *Y. lipolytica*, was expressed to evaluate BAK production under enhanced precursor-supply conditions (Fig. 3A). The resulting strain, YL04, produced BAK at a titer of 0.12 mg/L, as confirmed by shake-flask fermentation and HPLC analysis.

**Fig. 3.**
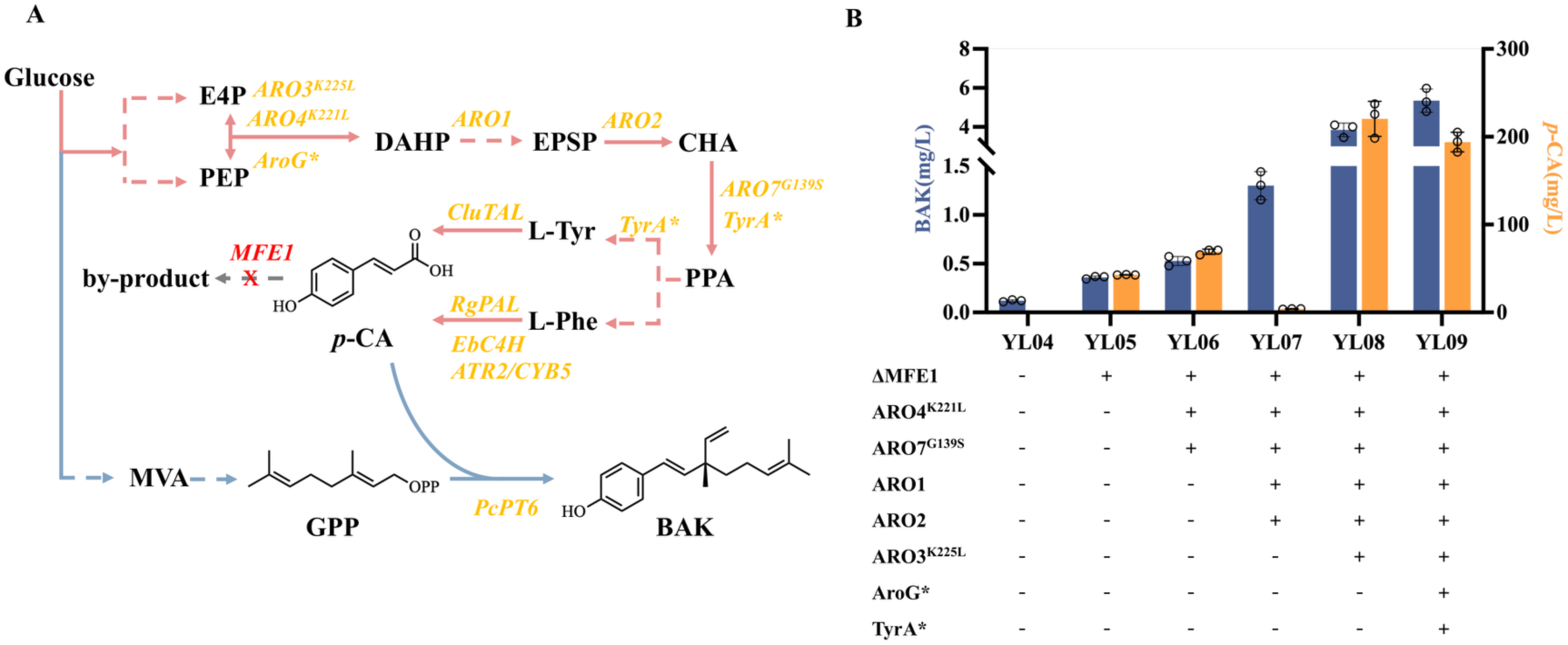
De novo synthesis and stepwise metabolic optimization of BAK in *Yarrowia lipolytica*. (A) Schematic of the reconstructed BAK biosynthetic pathway. The *p*-CA supply module includes *CluTAL*, *RgPAL*, *EbC4H*, *ATR2*, and *CYB5*, together with the feedback-resistant variants *ARO4*^K221L^, *ARO7*^G139S^, *ARO3*^K225L^, *AroG*\*, and *TyrA*\*. *ARO1* and *ARO2* were overexpressed to reinforce the shikimate pathway, and *MFE1* was deleted to reduce *p*-CA degradation. GPP was supplied through the MVA pathway, and PcPT6 catalyzed the terminal conversion of *p*-CA and GPP to BAK. Gene annotations in (A): *ARO3*^K225L^ and *ARO4*^K221L^ encoding 3-deoxy-D-arabino-heptulosonate 7-phosphate synthase variants; *ARO1* encoding pentafunctional AROM polypeptide; *ARO2* encoding chorismate synthase; *ARO7*^G139S^ encoding a feedback-resistant chorismate mutase variant; *AroG*\* encoding feedback-resistant 3-deoxy-D-arabino-heptulosonate 7-phosphate synthase from *Escherichia coli*; *TyrA*\* encoding feedback-resistant bifunctional chorismate mutase/prephenate dehydrogenase from *E. coli*; *CluTAL* encoding tyrosine ammonia-lyase; *RgPAL* encoding phenylalanine ammonia-lyase; *EbC4H* encoding cinnamate 4-hydroxylase; *ATR2* encoding NADPH-cytochrome P450 reductase 2 from *Arabidopsis thaliana*; *CYB5* encoding cytochrome b5 from *Y. lipolytica*; *MFE1* encoding peroxisomal beta-oxidation multifunctional enzyme; *PcPT6* encoding prenyltransferase 6 from *P. corylifolia*. K225L, K221L, and G139S denote Lys-to-Leu substitutions at residues 225 and 221 and a Gly-to-Ser substitution at residue 139, respectively; the asterisk denotes feedback-resistant variants. Abbreviations in (A): E4P, erythrose 4-phosphate; PEP, phosphoenolpyruvate; DAHP, 3-deoxy-D-arabino-heptulosonate 7-phosphate; EPSP, 5-enolpyruvylshikimate 3-phosphate; CHA, chorismate; PPA, prephenate; L-Tyr, L-tyrosine; L-Phe, L-phenylalanine; *p*-CA, *p*-coumaric acid; MVA, mevalonic acid; GPP, geranyl diphosphate; BAK, bakuchiol; OPP, diphosphate group in the GPP structure. (B) BAK production and residual *p*-CA accumulation in strains YL04-YL09 during stepwise pathway engineering. Data in (B) represent the mean of n = 3 biologically independent samples, and error bars show standard deviation (s.d.). Circles represent individual raw data points.

We next evaluated whether GPP availability limited BAK production. To enhance GPP supply, strain YL03 was constructed by overexpressing two key enzymes in the MVA pathway, truncated 3-hydroxy-3-methylglutaryl-CoA reductase (tHMGR) and isopentenyl pyrophosphate isomerase (IDI) (Yao et al., 2025), following previously reported strategies. However, under the same fermentation conditions, BAK production remained undetectable, and no additional product peaks were observed (Fig. S8). One possible explanation is that increased upstream isoprenoid flux altered the balance of prenyl diphosphate intermediates, such as GPP, FPP, and GGPP, with potential effects on MVA-pathway regulation (Primak et al., 2011). This explanation remains to be tested. The lack of detectable BAK nevertheless indicated that this intervention alone was insufficient in the YL00 background. We therefore focused subsequent engineering efforts on strengthening the *p*-CA biosynthetic module.

We subsequently performed a series of modifications targeting the shikimate and aromatic amino acid biosynthetic pathways and monitored both BAK production and *p*-CA accumulation in the resulting strains (Fig. 3B). First, *MFE1* was deleted in YL04 to reduce *p*-CA degradation (Zhu et al., 2026), generating YL05. This modification increased BAK production from 0.12 to 0.36 mg/L, reaching three times the original titer, while 42.84 mg/L *p*-CA accumulated in the culture. We next targeted feedback inhibition within the shikimate pathway by introducing the feedback-resistant mutants *ARO4*^K221L^ and *ARO7*^G139S^ into YL05 (Liu et al., 2025), generating YL06. BAK production increased modestly to 0.53 mg/L, representing a 47.2% increase relative to YL05, whereas *p*-CA accumulation further increased to 69.15 mg/L. The modest increase in BAK relative to *p*-CA accumulation suggested that relieving feedback inhibition at these steps was insufficient to remove the remaining constraints on BAK production.

We then overexpressed *ARO1* and *ARO2* (Wang *et al*., 2022) in YL06 to generate YL07. This modification substantially increased BAK production, while *p*-CA was no longer detectable in the culture. The combination of increased BAK production and undetectable residual *p*-CA was consistent with more complete precursor utilization. This response supported further engineering of aromatic precursor supply, although extracellular *p*-CA measurements alone do not establish intracellular flux or PcPT6 catalytic capacity.

To further increase aromatic precursor flux, the feedback-resistant mutant *ARO3*^K225L^ (Wang *et al*., 2022) was introduced into YL07, generating YL08. Compared with YL07, YL08 showed a pronounced increase in BAK production, reaching 3.85 mg/L, accompanied by a marked increase in *p*-CA accumulation to 220.44 mg/L. The simultaneous increase in both BAK production and residual *p*-CA supported a limitation in *p*-CA supply in YL07. Relieving feedback inhibition at the ARO3-catalyzed step therefore improved BAK production while increasing residual *p*-CA. Notably, introducing the feedback-resistant *ARO4* and *ARO7* mutants into YL05 resulted in only a modest improvement in BAK production, whereas introduction of the feedback-resistant *ARO3* mutant into YL07 produced a much greater increase. Because these modifications were tested in different strain backgrounds, the responses do not directly establish the relative contributions of ARO3, ARO4, and ARO7 to flux control. Finally, *AroG*\* and *TyrA*\* from *E. coli* (Zhang et al., 2023) were overexpressed in YL08 to further enhance DAHP formation and tyrosine biosynthesis, respectively, generating YL09. Compared with YL08, YL09 exhibited a 39.0% increase in BAK production, from 3.85 to 5.35 mg/L, whereas *p*-CA accumulation decreased by 12.0%, from 220.44 to 193.98 mg/L (Fig. 3C). The simultaneous increase in BAK production and reduction in residual *p*-CA suggested that enhancing DAHP and tyrosine biosynthesis favored BAK formation and improved *p*-CA utilization.

Together, these modifications increased BAK production by improving aromatic precursor supply through reduced *p*-CA degradation, relief of feedback inhibition, and reinforcement of shikimate and tyrosine biosynthesis. The changing relationship between BAK production and residual *p*-CA suggested that the main pathway constraints also changed during engineering. In particular, substantial *p*-CA accumulation in YL09 motivated further improvement of GPP supply and downstream precursor conversion using a coordinated push-pull strategy.

### 5. Coordinated push-pull engineering of dual precursors enhances BAK production

The substantial residual *p*-CA in YL09 suggested that additional gains in BAK production would require improved GPP supply, increased downstream conversion capacity, or both. We therefore adopted a push-pull strategy that combined reinforcement of the MVA pathway to increase GPP supply (push) with increased *t40PcPT6* expression to promote precursor conversion (pull) (Fig. 4A).

**Fig. 4.**
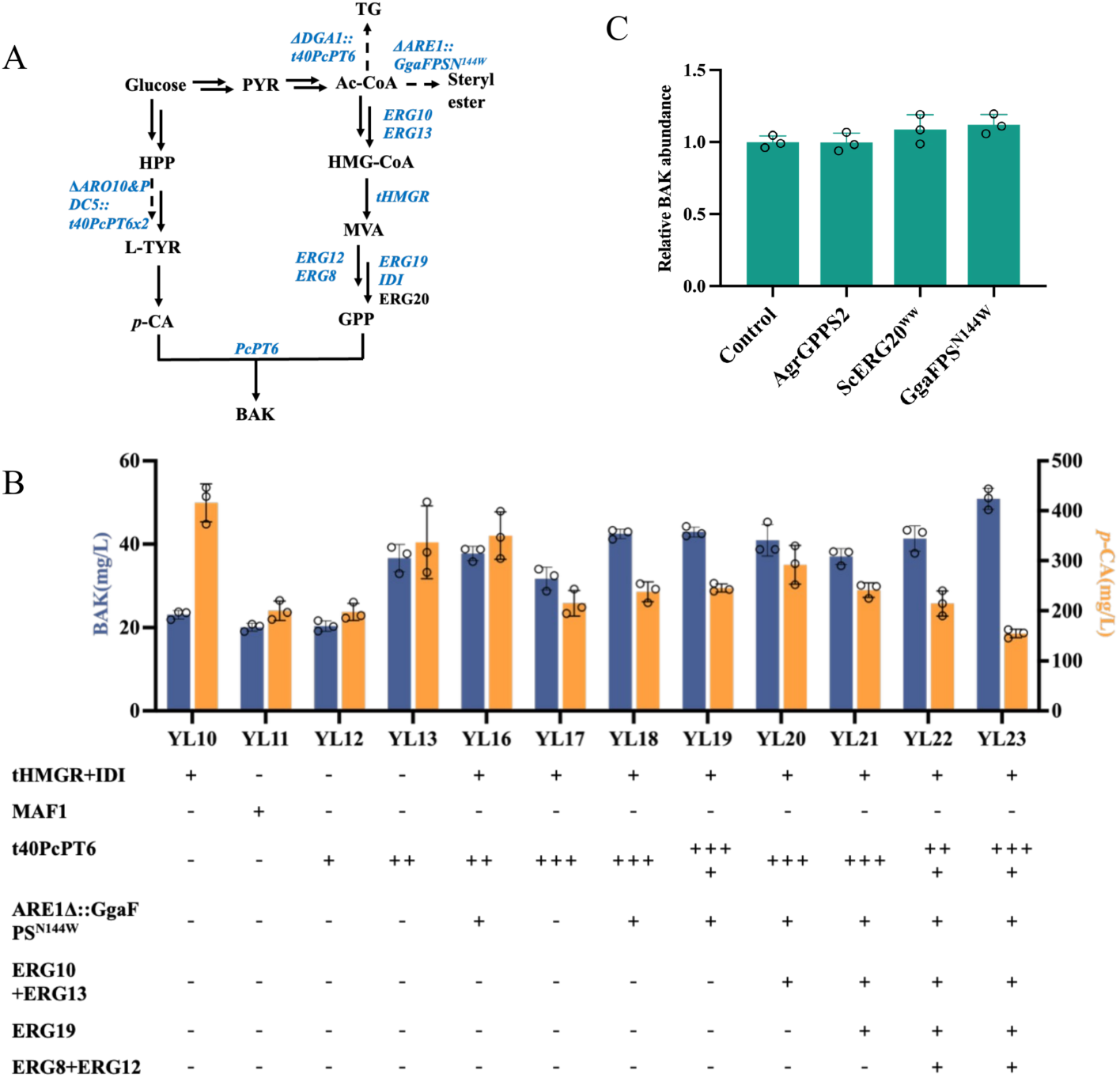
Coordinated “push-pull” strategy synergistically increases BAK production. (A) Schematic of the push-pull design. The MVA pathway was reinforced to increase GPP supply (push), whereas the copy number of *t40PcPT6* was progressively increased to enhance conversion of *p*-CA to BAK (pull). Deletion of *ARO10* and *PDC5* was used to redirect flux toward *p*-CA biosynthesis. Gene annotations in (A): *ERG10* encoding acetyl-CoA acetyltransferase (acetoacetyl-CoA thiolase); *ERG13* encoding 3-hydroxy-3-methylglutaryl-CoA synthase; *tHMGR* encoding truncated 3-hydroxy-3-methylglutaryl-CoA reductase; *ERG12* encoding mevalonate kinase; *ERG8* encoding phosphomevalonate kinase; *ERG19* encoding mevalonate diphosphate decarboxylase; *IDI* encoding isopentenyl diphosphate isomerase; *ERG20* encoding farnesyl diphosphate synthase; *ARO10* encoding phenylpyruvate decarboxylase; *PDC5* encoding pyruvate decarboxylase 5; *ARE1* encoding acyl-CoA:sterol O-acyltransferase; *DGA1* encoding acyl-CoA:diacylglycerol O-acyltransferase 1; *GgaFPS*^N144W^ encoding the N144W variant of farnesyl diphosphate synthase from *Gallus gallus*; *PcPT6* encoding prenyltransferase 6 from *P. corylifolia*; *t40PcPT6* encoding a PcPT6 variant lacking the first 40 N-terminal amino acids. N144W denotes an Asn-to-Trp substitution at residue 144. The symbol Δ indicates gene deletion,:: indicates insertion at the indicated genomic locus, and x2 indicates two copies. Abbreviations in (A): PYR, pyruvate; Ac-CoA, acetyl-coenzyme A; HMG-CoA, 3-hydroxy-3-methylglutaryl-coenzyme A; HPP, 4-hydroxyphenylpyruvate; L-TYR, L-tyrosine; *p*-CA, *p*-coumaric acid; MVA, mevalonic acid; GPP, geranyl diphosphate; BAK, bakuchiol; TG, triacylglycerol. (B) BAK production and residual *p*-CA in strains YL10-YL23. Data in (B, C) represent the mean of n = 3 biologically independent samples, and error bars show standard deviation (s.d.). Circles represent individual raw data points.

We first evaluated two strategies for enhancing GPP supply. In strain YL09, overexpression of *tHMGR* and *IDI* generated strain YL10, with the aim of increasing GPP supply. YL10 produced 23.08 mg/L BAK, corresponding to 4.31 times the titer of YL09, while *p*-CA accumulation reached 2.14 times the level in YL09, at 415.90 mg/L. As an alternative approach, MAF1, a highly conserved negative regulator of RNA polymerase III, was overexpressed to potentially increase the intracellular DMAPP pool and thereby promote GPP formation (Liu et al., 2013b). The resulting strain YL11 produced slightly less BAK than YL10. Therefore, YL10 was selected as the starting strain for subsequent engineering. Under the tested conditions, *tHMGR* and *IDI* overexpression improved BAK production more than *MAF1* overexpression.

We next sought to further improve precursor availability by redirecting metabolic flux toward *p*-CA biosynthesis. In YL10, *ARO10* and *PDC5* were deleted (Liu et al., 2026; Xin et al., 2025), and *t40PcPT6* was integrated at the corresponding loci to generate strain YL13 carrying two additional copies of *t40PcPT6*. Compared with YL10, YL13 exhibited a 58.62% increase in BAK production, from 23.08 to 36.61 mg/L, whereas *p*-CA production decreased by 19.0%, from 415.90 to approximately 337.06 mg/L. The increase in BAK production and decrease in residual *p*-CA were consistent with improved precursor utilization. Because gene deletions and *t40PcPT6* integration were introduced together, their individual contributions could not be separated in this comparison.

We then investigated whether further increasing GPP availability could improve BAK production. Introduction of the GgaFPS^N144W^ GPP synthase variant from *Gallus gallus* (Brown et al., 2015) into YL13 generated strain YL16, resulting in a 3.11% increase in BAK production relative to YL13. However, the relatively modest improvement suggested that this intervention alone had a limited effect at this stage. We therefore further examined the effect of increasing the *t40PcPT6* copy number. Increasing *t40PcPT6* to three copies in the absence of GgaFPS^N144W^ generated YL17 and led to a pronounced decrease in BAK production, suggesting that the effect of additional *t40PcPT6* copies depended on the precursor-supply background. In contrast, simultaneous expression of GgaFPS^N144W^ and increasing the *t40PcPT6* copy number to three generated YL18, which produced 1.13 times as much BAK as YL16 and exhibited a concomitant decrease in *p*-CA accumulation. The improved performance of YL18 supported coordinated engineering of GPP supply and precursor conversion.

To further strengthen this strategy, we enhanced the upstream MVA pathway by overexpressing *ERG10, ERG13, ERG19, ERG8, and ERG12* (Zhang et al., 2024; Zhang et al., 2025), while simultaneously increasing the *t40PcPT6* copy number to four, generating strain YL23. Compared with YL18, YL23 showed a 19.81% increase in BAK production, from 42.46 to 50.87 mg/L, accompanied by a 36.97% reduction in *p*-CA accumulation, from 245.79 to 154.92 mg/L. In contrast, increasing the *t40PcPT6* copy number to four in the absence of further MVA pathway reinforcement (YL19) resulted in only marginal changes, with BAK production and *p*-CA accumulation changing by 1.04% and 3.25%, respectively. These comparisons further supported reinforcing GPP supply alongside *t40PcPT6* expression. Accordingly, strain YL23, which exhibited the highest BAK production among the engineered strains, was selected for subsequent fermentation optimization.

### 6. High-level production of BAK in fed-batch fermentation

Culture medium composition has been reported to be a critical determinant of metabolite production in *Y. lipolytica* (Jacobsen et al., 2020; Sáez-Sáez et al., 2020).

We therefore evaluated seven media with different nutritional compositions (CM, YPD, YNB, MM, MMY, MMP, and MMYP) for BAK production by strain YL23 (Fig. 5A). Among the media tested, MMY, consisting of minimal medium supplemented with yeast extract, supported the highest BAK production, reaching 108.29 mg/L, corresponding to 2.12 times the titer obtained in CM. In contrast, MMYP, supplemented with both yeast extract and peptone, supported the highest biomass accumulation, approximately 2.5 times that in CM, but yielded the lowest BAK titer, corresponding to only 78.56% of that in CM. These results indicate that enhanced biomass accumulation does not necessarily translate into increased BAK production. Yeast extract may provide vitamins or other growth factors that support BAK biosynthesis, whereas the additional nutrients in MMYP may favor biomass formation. These possible explanations require further investigation. Based on these results, MMY was selected for subsequent fed-batch fermentation.

**Fig. 5.**
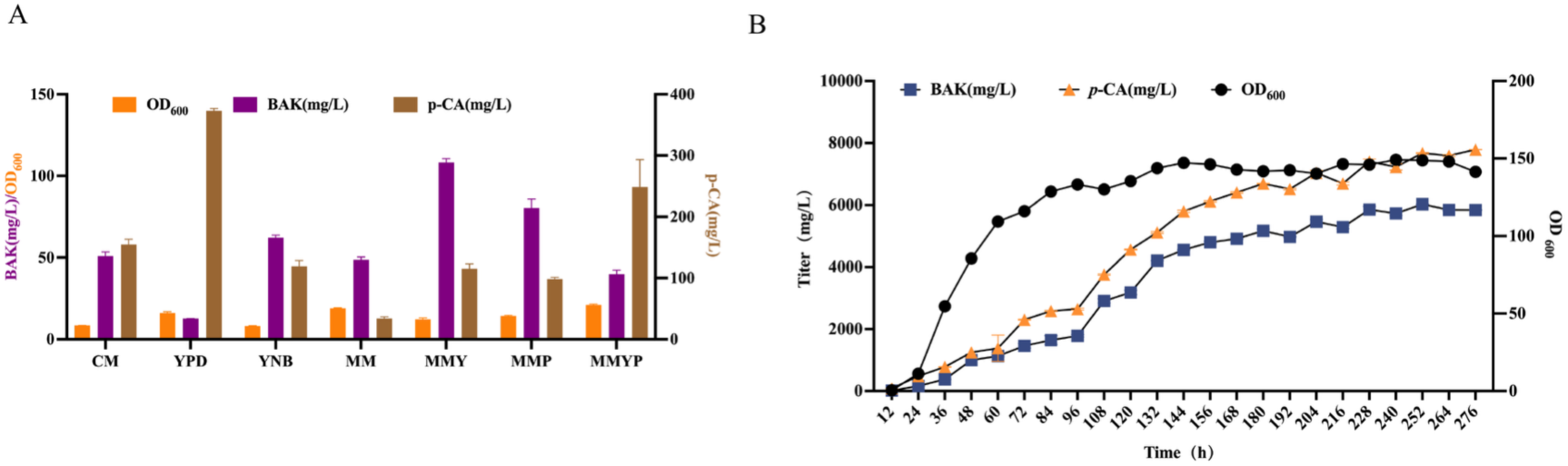
High-level production of BAK in a 7.5-L fed-batch fermentation. (A) Culture-medium optimization for strain YL23. Seven media (CM, YPD, YNB, MM, MMY, MMP, and MMYP) were compared for BAK titer, *p*-CA accumulation, OD600, and BAK titer normalized to OD600. MMY produced the highest BAK titer (108.29 mg/L, 2.12-fold that in CM). (B) Time-course profiles of BAK titer, residual *p*-CA, and OD600 during fed-batch fermentation in a 7.5-L bioreactor. BAK reached 6,026.57 mg/L at 252 h, with a maximum volumetric productivity of 643 mg/L/d between 96 and 108 h. Corn steep powder was omitted from the initial medium and supplied intermittently at 144, 168, 192, and 216 h, and the pH was adjusted to 6.5 after 96 h. At the maximum BAK titer, residual *p*-CA reached 7,674.11 mg/L. Data in (A) represent the mean of n = 3 biologically independent samples, and error bars show standard deviation (s.d.).

To enhance BAK production under high-cell-density conditions, fed-batch fermentation was performed in a 7.5-L bioreactor using MMY. To avoid excessive biomass accumulation during the later stages of fermentation, corn steep powder was excluded from the initial medium and supplemented intermittently at 144, 168, 192, and 216 h. The pH was adjusted to 6.5 after 96 h to stabilize the fermentation conditions. BAK production increased steadily and reached a maximum titer of approximately 6.0 g/L at 252 h (Fig. 5B, S9), achieving gram-per-liter BAK production. Notably, BAK accumulation continued after biomass growth had begun to plateau, indicating that product formation was partially uncoupled from cell growth. The highest volumetric BAK productivity, 643 mg/L/day, was observed between 96 and 108 h, coinciding with a period in which biomass accumulation had begun to slow, consistent with continued product formation as growth slowed. At the maximum BAK titer, *p*-CA had accumulated to 7,674.11 mg/L. The substantial residual *p*-CA suggests that *p*-CA conversion, rather than its biosynthetic supply, may represent a major constraint on BAK production. Possible constraints include GPP availability, PcPT6 expression or catalytic capacity, and substrate accessibility. Following the peak in BAK production, the BAK titer decreased slightly, potentially due to product degradation or further metabolism.

## Discussion

This study connects a chromosome-level *P. corylifolia* genome with functional screening of legume PTs and microbial BAK production. The assembly’s contiguity and completeness support systematic gene identification, while comparative analyses place these candidates in their genomic and evolutionary context. The sister relationship with soybean and retention of the ancestral legume duplication signature provide a framework for examining metabolic gene diversification. The abundance of LTR retrotransposons also warrants investigation of their contribution to genome structure. These observations provide context for future functional studies, but do not establish the evolutionary origin of BAK-producing activity. Beyond BAK biosynthesis, the genome supports investigation of other specialized metabolic pathways in *P. corylifolia*. Genome-wide identification broadened the PT candidate pool, although tissue expression and structural predictions were subsequently used to prioritize experimental screening. PcPT6 corresponds to the previously reported PcPT07 (Zheng et al., 2024), so its activity confirms an established catalytic function. The additional active *P. corylifolia* and *V. radiata* homologs extend the experimentally supported sequence and species range of PTs capable of supporting BAK production in yeast. Activity of the truncated *V. radiata* enzymes does not, however, establish their native physiological roles or pathway contributions in the plant.

PcPT6 also accepted selected hydroxycinnamic acid analogues, providing a route to BAK derivatives relevant to studies of their reported biological activities (Galvez et al., 2026). Caffeic acid yielded 3-hydroxybakuchiol, whereas the product from 2,4-dihydroxycinnamic acid remains putatively assigned as 2-hydroxybakuchiol (Fig. S7C, D). The latter assignment requires confirmation because an authentic standard was unavailable. The absence of detectable conversion of ferulic acid suggests constraints on substrate recognition or productive positioning. Its 3-methoxy group is a possible contributor, but the current substrate panel does not establish a steric mechanism.

N-terminal truncation provided a practical means of improving PcPT6 performance in yeast. Removal of 40 amino acids increased BAK production by 53%, whereas more extensive truncation sharply reduced production. This restricted window suggests that the N-terminal region influences heterologous enzyme performance while remaining coupled to protein integrity or membrane association. Because activity was assessed through whole-cell product formation, the improvement cannot be assigned specifically to intrinsic catalytic efficiency. Differences in expression, stability, localization and substrate accessibility remain plausible contributors. Matched measurements of enzyme abundance and membrane-associated activity would distinguish these explanations and guide further engineering.

The pathway-engineering results support coordinating precursor supply with terminal conversion rather than increasing either component in isolation. Reinforcing aromatic precursor synthesis increased BAK production but also exposed a persistent excess of residual *p*-CA. Additional *t40PcPT6* copies produced different responses across GPP-supply backgrounds, showing that gene dosage alone did not predict production performance. Reinforcement of the MVA pathway alongside increased PT gene dosage supported further improvement. These comparisons identify a useful engineering strategy, but extracellular metabolite concentrations do not quantify intracellular flux. Limited GPP availability, expression burden and membrane-associated constraints could each contribute to the observed responses.

Fermentation optimization translated pathway engineering into gram-per-liter BAK production, reaching 6.03 g/L in a 7.5-L fed-batch bioreactor. The contrast between biomass accumulation and BAK production across media indicates that conditions favoring growth were not necessarily optimal for product formation. Continued BAK accumulation as biomass growth slowed is consistent with partial decoupling of production from active proliferation. Residual glucose during late fermentation argues against extracellular glucose depletion as the sole limitation. However, nutrient supply remained important, as reducing the feeding rate decreased both biomass and BAK production. Reported BAK contents of 1-3% of *P. corylifolia* seed dry weight (Adarsh Krishna *et al*., 2022) cannot be compared directly with fermentation titers. A common basis for yield, productivity and product recovery would be needed to evaluate the relative performance of plant extraction and microbial production.

Persistent *p*-CA accumulation and the decline in BAK titer after its maximum define two priorities for further optimization. Measurements of GPP availability and PT abundance would help separate precursor limitation from constraints on terminal conversion. Dynamic pathway regulation could then be tested against the specific imbalance identified, rather than assuming that additional constitutive expression will improve production. In parallel, tracing BAK during prolonged cultivation would distinguish degradation or further metabolism from changes in product recovery. These tests would address the remaining production constraints while preserving the demonstrated advantages of coordinated pathway engineering.

## Conclusion

This study establishes a chromosome-level genomic resource for P. corylifolia and extends functional evidence for BAK-producing PTs to homologs from P. corylifolia and V. radiata. N-terminal enzyme engineering, coordinated precursor supply and terminal conversion, and fermentation optimization together supported 6.03 g/L BAK production in Y. lipolytica. The resulting genomic resource, validated enzymes and production platform provide a basis for further development of microbial BAK biosynthesis. Resolving the remaining limits on precursor conversion and product retention will be important for improving process yield and recovery.

## Materials and methods

### DNA and RNA isolation and sequencing

*P. corylifolia* used for genome sequencing was collected from the Institute of Tropical Eco-Agriculture, Yunnan Academy of Agricultural Sciences, China (25.70°N, 101.87°E). Genomic DNA was extracted from young leaves of *P. corylifolia* using an improved cetyltrimethylammonium bromide (CTAB) method. Genome sequencing was performed using both PacBio long-read sequencing and Illumina high-throughput sequencing technologies (Benagen LTD). Total plant RNA was extracted using the RNAprep Pure Plant Kit (Tiangen, Beijing, China). RNA concentration, purity, and integrity were evaluated using a NanoDrop 2000 (Thermo Fisher Scientific) and an Agilent Bioanalyzer 2100 system (Agilent Technologies). Enriched mRNA libraries were constructed from 1 μg of total RNA using the Hieff NGS Ultima Dual-Mode mRNA Library Prep Kit (Yeasen Biotechnology, Shanghai, China), which involved poly-T magnetic bead purification, fragmentation, cDNA synthesis, end-repair, 3′ adenylation, adapter ligation, and PCR enrichment. After purification via the AMPure XP system (Beckman Coulter) and quality assessment, the libraries were sequenced on an Illumina NovaSeq platform to yield 150-bp paired-end reads.

### Estimation of *P. corylifolia* genome size

To estimate the genomic characteristics of *P. corylifolia*, raw short reads from the Illumina platform were first quality-filtered using fastp (v0.23.4) with the following parameters:-q 10-u 50-y-g-Y 10-e 20-l 100-b 150-B 150 to remove low-quality bases and short adapter-contaminated reads (Chen et al., 2018). The high-quality filtered reads were subsequently subjected to K-mer frequency distribution analysis (k = 21) using Jellyfish software (Marcais and Kingsford, 2011). Based on the K-mer depth distribution, the genome size and genomic traits were calculated using the GCE pipeline (Liu et al., 2013a). The genome size of *P. corylifolia* was estimated to be approximately 484.06 Mb, characterized by a repetitive sequence content of 51.03% and a low heterozygosity rate of 0.03%.

### Genome assembly and chromosome-level scaffolding

The Illumina sequencing platform was used to generate 55.72 Gb of raw sequences with a sequencing depth of about 125.41× for estimated genome 484.06 Mb and a GC content of 34.39%. Using Hifiasm (v0.19), the draft genome of *P. corylifolia* was assembled from 39.3 Gb of PacBio HiFi clean reads (reads N50: 15.14Kb; mean length: 14.76 Kb) (Cheng et al., 2020). The initial assembly comprised 504.65 Mb of contig sequences with a contig N50 of 40.33 Mb. Foreign contaminants and mitochondrial and plastid sequences were identified by alignment against the NCBI NT and organelle databases and removed from the assembly.

For scaffolding, 100.86 Gb of Hi-C data were aligned to the polished assembly using BWA (v0.7.17) under default parameters (Li and Durbin, 2009). After filtering invalid reads (e.g., self-ligations, non-ligations, PCR duplicates, and anomalous fragments), valid paired-end reads mapping to different contigs were used for scaffolding. Utilizing the agglomerative hierarchical clustering in LACHESIS, 69 contigs were clustered into 63 groups, ordered, and oriented (Burton et al., 2013). In the final chromosome-level assembly, approximately 471.61 Mb of sequence was anchored to 11 chromosomes, accounting for 99.97% of the total assembly. Of the sequences mapped onto chromosomes, approximately 467.75 Mb had a defined order and orientation, representing 99.18% of the total length of the mapped chromosomal sequences. Assembly completeness was validated via BUSCO (v5.2.2) using the embryophyta_odb10 dataset (1,614 groups) (Simao et al., 2015).

### Annotation of protein-coding genes and repetitive sequences

The protein-coding gene models of *P. corylifolia* were predicted using an integrated pipeline combining *ab initio* prediction, homology search, and transcript-based assembly. Specifically, Augustus (v3.1.0) was used for *ab initio* gene modeling (Stanke et al., 2008). GeMoMa (v1.7) was applied for homology-based prediction using reference gene sets from closely related species (Keilwagen et al., 2016). For transcript-based annotation, RNA-seq reads were mapped onto the genome assembly via HISAT2 (v2.1.0) and assembled using StringTie (v2.1.4), with subsequent gene prediction performed by GeneMarkS-T (v5.1) (Kim et al., 2015; Pertea et al., 2015; Tang et al., 2015). Furthermore, PASA (v2.4.1) was used to incorporate unigenes and full-length transcripts assembled by Trinity (v2.11) (Grabherr et al., 2011; Haas et al., 2003). Finally, EVidenceModeler (v1.1.1) was employed to integrate the gene models, which were then refined using PASA (Haas et al., 2008). This pipeline yielded 31,907 protein-coding genes (average length: 3,713.69 bp), with 99.18% successfully localized on the 11 pseudochromosomes. Annotation completeness was evaluated using BUSCO (v5.2.2) (Simao *et al*., 2015).

Repetitive elements, including transposable elements (TEs) and tandem repeats, were annotated. A species-specific *de novo* repeat library was generated using RepeatModeler (v2.0.1) via its internal RECON (Bao and Eddy, 2002) and RepeatScout algorithms (Flynn et al., 2020; Price et al., 2005). Structurally intact full-length long terminal repeat retrotransposons (fl-LTR-RTs) were detected using LTRharvest (v1.5.9) and LTR_FINDER (v2.8) and further refined into a non-redundant LTR library using LTR_retriever (Ellinghaus et al., 2008; Ou and Jiang, 2018; Xu and Wang, 2007). By combining these de novo predictions with the known Dfam (v3.5) database, a final non-redundant TE library was constructed. RepeatMasker (v4.1.2) was then utilized to identify and classify genomic TEs based on this combined library (Tarailo-Graovac and Chen, 2009). In parallel, tandem repeats and microsatellites were identified using Tandem Repeats Finder (TRF, v4.09) and MISA (v2.1), respectively (Beier et al., 2017; Benson, 1999).

### Functional annotation of protein-coding genes

For functional annotation, the protein sequences encoded by the 31,907 predicted genes were aligned against public databases, including the NCBI Non-Redundant protein (NR), eggNOG, Clusters of Orthologous Groups of proteins (KOG), TrEMBL, and Swiss-Prot (manually annotated and reviewed) databases, as well as the Kyoto Encyclopedia of Genes and Genomes (KEGG) Ortholog (KO) database. These alignments were performed using DIAMOND blastp (v0.9.29.130) with an *E*-value cutoff of 1×10^-3^. Additionally, InterProScan (v5.34-73.0) and the Pfam (Protein family) database were employed to identify conserved protein domains and motifs (Jones et al., 2014; Mistry et al., 2021). Gene Ontology (GO) terms for individual genes were subsequently integrated from TrEMBL, InterPro, and eggNOG annotation tracks (Huerta-Cepas et al., 2019; Yip et al., 2004). In total, 99.02% of the entire gene set was assigned functional descriptions, conserved domains, or GO terms.

### Annotation of non-coding RNA genes

Non-coding RNA (ncRNA) genes in the *P. corylifolia* genome were annotated using dedicated computational tools. Transfer RNAs (tRNAs) were predicted using tRNAscan-SE (v1.3.1) with default parameters (Lowe and Eddy, 1997). Ribosomal RNAs (rRNAs) were identified using Barrnap (v0.9) under default settings. Other regulatory small non-coding RNAs, including microRNAs (miRNAs), small nuclear RNAs (snRNAs), and small nucleolar RNAs (snoRNAs), were identified and classified using Infernal (v1.1) by searching against the Rfam database (v14.5) with default parameters (Griffiths-Jones et al., 2005; Nawrocki and Eddy, 2013).

### Phylogenetic tree construction and divergence time estimation

To resolve the phylogenetic relationships of *P. corylifolia* within the Fabaceae family, orthologous gene families across 16 species were identified using OrthoFinder (v2.5.5) with default parameters (Emms and Kelly, 2019). A total of 117 single-copy orthologs shared by all 16 genomes, including *Glycine max* and other representative legumes, were selected for phylogenetic reconstruction. Protein sequences for each ortholog were individually aligned using MAFFT (v7.490) and subsequently concatenated into a sequence supermatrix (Rozewicki et al., 2019). A maximum-likelihood phylogenetic tree was then constructed using IQ-TREE (v1.6.12) (Minh et al., 2020). Based on this topology, divergence times were estimated under a relaxed clock model using the r8s program with default parameters(Sanderson, 2003). Calibration points and fossil-based time constraints were retrieved from the TimeTree database (http://www.timetree.org/) (Kumar et al., 2022). The final chronogram was visualized and refined using custom Python scripts and Interactive Tree of Life (iTOL) (Letunic and Bork, 2024).

### Gene family expansion and contraction, synteny, and whole-genome duplication analysis

Gene family expansion and contraction dynamics across the 16 sequenced species were analyzed using CAFE5 (Mendes et al., 2020). The phylogenetic tree topology and branch lengths were used to infer the statistical significance of changes in gene family size along each lineage. For comparative genomics, a genome-wide synteny analysis among *P. corylifolia, C. cajan, P. vulgaris, V. radiata, and G. max* was performed using the JCVI toolkit (MCScan Python version 1.5.4) based on genomic annotation (GFF3) files (Tang et al., 2024). To investigate whole-genome duplication events, the synonymous substitution rates (*Ks*) of the identified syntenic gene pairs were calculated using the Whole-Genome Duplication Identifier (WGDI) pipeline (v0.7.4) (Sun et al., 2022).

### Gene expression quantification and differential expression analysis

Gene expression was quantified as fragments per kilobase of transcript per million mapped fragments (FPKM). RNA-seq data collected from four distinct *P. corylifolia* tissues (root, stem, leaf, and seed), each represented by three biological replicates, were aligned to the reference genome using HISAT2 (Kim *et al*., 2015). Differential expression analysis between tissues or groups was performed using DESeq2 (Roy et al., 2026). The resulting *P*-values were adjusted using the Benjamini-Hochberg approach to control the false discovery rate (FDR). Genes with adjusted *P*-values < 0.01 and |log2FC| ≥1 were classified as differentially expressed genes (DEGs).

### Identification and screening of prenyltransferases

To identify candidate prenyltransferases, a genome-wide search was performed against the annotated *P. corylifolia* genome database using hmmscan (HMMER v3.3.1) (Mistry *et al*., 2021; Potter et al., 2018). Protein sequences containing the prenyltransferase Pfam domain (PF01040) were retrieved with an *E*-value threshold of 1×10^-4^. For evolutionary analysis, multiple sequence alignment of the identified PTs was conducted using Clustal Omega (v1.2.4) under default parameters (Sievers and Higgins, 2018). A maximum-likelihood (ML) phylogenetic tree was subsequently constructed using RAxML (v8.2.12, PTHREADS-SSE3 version) under the PROTGAMMALGX amino acid substitution model with 100 bootstrap replicates(Stamatakis, 2014). The finalized phylogenetic tree was visualized and annotated using iTOL. Furthermore, the three-dimensional protein structures of the candidate enzymes were predicted using AlphaFold3 to evaluate their structural characteristics (Jumper et al., 2021).

### Plant materials, reagents, media, and culture conditions

The plant material *Psoralea corylifolia* was collected from the Hot Region Ecological Agricultural Research Institute, Yunnan Academy of Agricultural Sciences. Both bakuchiol and *p*-coumaric acid were of ultra-pure grade. Corn steep powder was obtained from HONGRUNBAOSHU (China). All other chemicals were of analytical grade unless otherwise indicated. *Escherichia coli* cells were cultured in LB medium containing 100 mg/L ampicillin, at 37°C and 220 rpm. The wild-type *Saccharomyces cerevisiae strain* W303-1B and *Y. lipolytica* strain W29 (ATCC No. 20460) were cultured in YPD or CM medium at 30°C and 220 rpm. The gRNA plasmid pCfB6627 was obtained from Addgene (https://www.addgene.org/), plasmid #106159. MMY medium for fed-batch fermentation contained 7.5 g/L ammonium sulfate, 14.4 g/L potassium dihydrogen phosphate, 0.5 g/L magnesium sulfate heptahydrate, 40 g/L glucose, and 10 g/L yeast extract. The feed medium contained 600 g/L glucose and 2.5% (v/v) polysorbate 80.

### Construction of plasmids and strains

All primers were synthesized by Tsingke Biotechnology Co. (China). All codon-optimized heterologous genes were synthesized by GenScript (China). Recombinant vectors were constructed by Gibson Assembly using the super fusion cloning mix (US Everbright). All primers, plasmids, strains, and genes used in this study are listed in Supplementary information.

Transformation of *Saccharomyces cerevisiae* (Liu et al., 2018) and *Yarrowia lipolytica* was performed using previously reported lithium acetate methods (Holkenbrink et al., 2018; Wang *et al*., 2022).

### Functional characterization of prenyltransferases

To construct *S. cerevisiae* strains for screening prenyltransferases, plasmid Y33-ScERG20^ww^ was co-transformed into W303-1B with one of the prenyltransferase plasmids Y22-PcPT1 to Y22-PcPT6, Y22-t40PcPT6, or Y22-tVrPT1 to Y22-tVrPT3 using the lithium acetate method. Positive transformants were screened on CM medium lacking tryptophan and uracil (CM-Trp-Ura). After 48 hours, single colonies were picked from the plates and transferred to tubes containing 4 mL of CM-Trp-Ura medium. The culture was then grown overnight at 30°C and 220 rpm. The seed culture was inoculated at a 1:50 ratio into 50 mL of CM-Trp-Ura medium in an Erlenmeyer flask. After 12-14 hours of fermentation, the substrate *p*-CA was added to a final concentration of 1.5 mM, and fermentation continued for 72 hours. After fermentation, 50 mL of ethyl acetate was added to the shake flask, and the mixture was extracted for one hour and then centrifuged at 5000 rpm for 15 minutes to separate the aqueous and organic phases. The upper organic phase was collected and concentrated by rotary evaporation. The residue was reconstituted in 200 μL of methanol for HPLC and LC/MS analysis.

### Substrate range of PcPT6

Single colonies were picked from the plates and transferred to test tubes containing 4 mL of sterile CM medium lacking tryptophan and uracil. The seed culture was inoculated at a 1:50 ratio into an Erlenmeyer flask containing 50 mL of CM medium lacking tryptophan and uracil. After fermentation at 30°C and 200 rpm for 12-14 hours, the following substrates were added in separate cultures: 3-(4-hydroxyphenyl) propionic acid, cinnamic acid, 4-methylcinnamic acid, 3-hydroxycinnamic acid, 2,4-dihydroxycinnamic acid, caffeic acid, ferulic acid, dihydrocaffeic acid, 3,4-dimethoxyphenylpropionic acid, or coumaric acid. Each substrate was added to a final concentration of 1.5 mM. Fermentation was continued for 72 hours before extraction and analysis by HPLC and LC/MS.

### Detection of bakuchiol in legume seeds

Seeds from each legume species (10 g) were ground to a powder. The powder was mixed with 15 mL of methanol/acetone (1:1) and subjected to ultrasonic extraction for 30 min. The mixture was centrifuged at 12,000 rpm for 30 min, and the supernatant was collected for LC-MS analysis.

### Shake-flask fermentation and BAK quantification in recombinant *Y. lipolytica*

Recombinant BAK-producing *Y. lipolytica* strains were streaked onto YPD plates. After overnight culture, a single colony was picked and inoculated into 5 mL of YPD medium for 12-16 hours. The seed culture was then inoculated into a shake flask containing 30 mL of CM medium at a ratio of 1:30. Fermentation was performed at 30°C and 220 rpm for 4 days. The product was then extracted with methanol and quantified by HPLC.

### HPLC and mass spectrometry analysis

HPLC analysis was performed on a Shimadzu LC-20ADXR HPLC system equipped with a photodiode array (PDA) detector. Separation was achieved on a Waters C18 reversed-phase column (4.6 × 250 mm, 5 μm particle size) at a flow rate of 0.6 mL/min. The detection wavelength was set at 260 nm. The mobile phase consisted of solvent A (water containing 0.1% formic acid) and solvent B (acetonitrile). The gradient elution program was as follows: 10 min, 75% B; 15 min, 90% B; 20 min, 97% B; 25 min, 97% B; 27 min, 10% B; 30 min, 10% B.

Mass spectrometric analysis was performed on a Bruker micrOTOF-II mass spectrometer equipped with an electrospray ionization (ESI) source. Data were acquired in negative ion mode over an m/z range of 50-1000. The nitrogen flow rate was 6.0 L/min, the source temperature was 180 °C, the nebulizer pressure was 1 bar, and the probe voltage was 14.5 kV.

### Fed-batch fermentation

A single colony was inoculated into 5 mL of YPD medium and cultured for approximately 24 h. Subsequently, 1 mL of this primary culture was transferred to a 250 mL flask containing 50 mL of YPD medium to prepare the seed culture, which was incubated for an additional 24 h. The resulting seed culture was then inoculated at 5% (v/v) into a 7.5 L bioreactor (Xan-Bio, China) containing 2 L of MMY medium. Fermentation was conducted at 30 °C, and the pH was controlled by automatic ammonia addition at 5.5 for the first 96 h and at 6.5 thereafter. The aeration rate was set to 2.5 vvm, and dissolved oxygen (DO) was maintained above 40% saturation using an agitation cascade (200-900 rpm). A feed medium containing 600 g/L glucose and 2.5% (v/v) polysorbate 80 was introduced when the initial glucose in the bioreactor was depleted. The feeding rate was adjusted based on biomass levels, ranging from 10 to 25 mL/h. A suspension containing 5 g of corn steep powder was added at 144, 168, 192, and 216 h. Cell growth and the concentrations of residual glucose, *p*-CA, and BAK were monitored every 12 h throughout the fermentation process.

### Measurement of OD600 and residual glucose

Cell growth was assessed by measuring the absorbance of a 1 mL fermentation broth sample at 600 nm using a UV spectrophotometer (AOELAB, China). A separate 1 mL sample was centrifuged at 12,000 rpm for 10 min, and the supernatant was collected for residual glucose measurement using a biochemical analyzer (Jinan Yanhe Biotechnology Co., Ltd.).

### Statistics and reproducibility

All experiments were performed with at least three biological replicates (n ≥ 3 per group), with error bars indicating standard deviation (s.d.). Data were analyzed using Excel or GraphPad Prism.

## Data availability

The data reported in this article are available in the main text or the Supplementary Information. The genome of *Senna tora, Abrus pulchellus, Arachis hypogaea, Sophora favescens, Cicer arietinum, Medicago sativa, Glycyrrhiza uralensis, Glycine max, Pueraria lobata, Phaseolus vulgaris, Vicia faba, Vigna radiata, Vigna angularis, Vigna unguiculata, Vigna mungo*, *Vigna reflexopilosa*, *Vigna umbellata*, *V. subterranea Cajanus cajan, Gastrolobium bilobum, Lotus japonicus, Medicago truncatula,* and *Mucuna pruriens* were Fetched in NCBI (National Center for Biotechnology Information). The genome data of this study are available from the corresponding author upon reasonable request.

## Author Contributions

Jing Li: designed and wrote the paper and conducted relevant experiments. Hong Chang: genome assembly and bioinformatics analysis. Chuan Liu: fed-batch fermentation. Shuai Li: Investigation. Jin Zhang: Investigation. Jie Bai: Investigation. Han Wang: Investigation.. Xiaonan Liu: conductd the relevant experiments and revised the manuscript. Zhijin Fan: revised the manuscript. Huifeng Jiang: revised the manuscript.

## Notes

The authors declare no competing financial interest.

## Acknowledgments

This work was supported by the National Natural Science Foundation of China (32472603, 32371499), Joint Fund Project of the National Natural Science Foundation of China (U23A20219), the National Key 542 Research & Development Program of China [2022YFD1700400 and 2022YFD1700402], Key Science & Technology Project of Anhui Province (202523n10050003, 202423l10050046), Youth Science and Technology Talent Cultivation Special Program of Hubei Province (2025DJA024), the Science and Technology Major Project of Guangxi (Guike AA24206050-3, Guike 2025JF2504850012, Guike AA24206048), Hubei University of Technology High-Level Talent Research Startup Fund Program (4301/00960), Hubei Provincial Key R&D Major Projects (2025BCA006), Natural Science Foundation of Wuhan (2025040601020158).

