## Supplementary information for "Chromosome-level genomics and cross-species enzyme screening enable high-titer bakuchiol production in *Yarrowia lipolytica*"

### Supplementary Figures

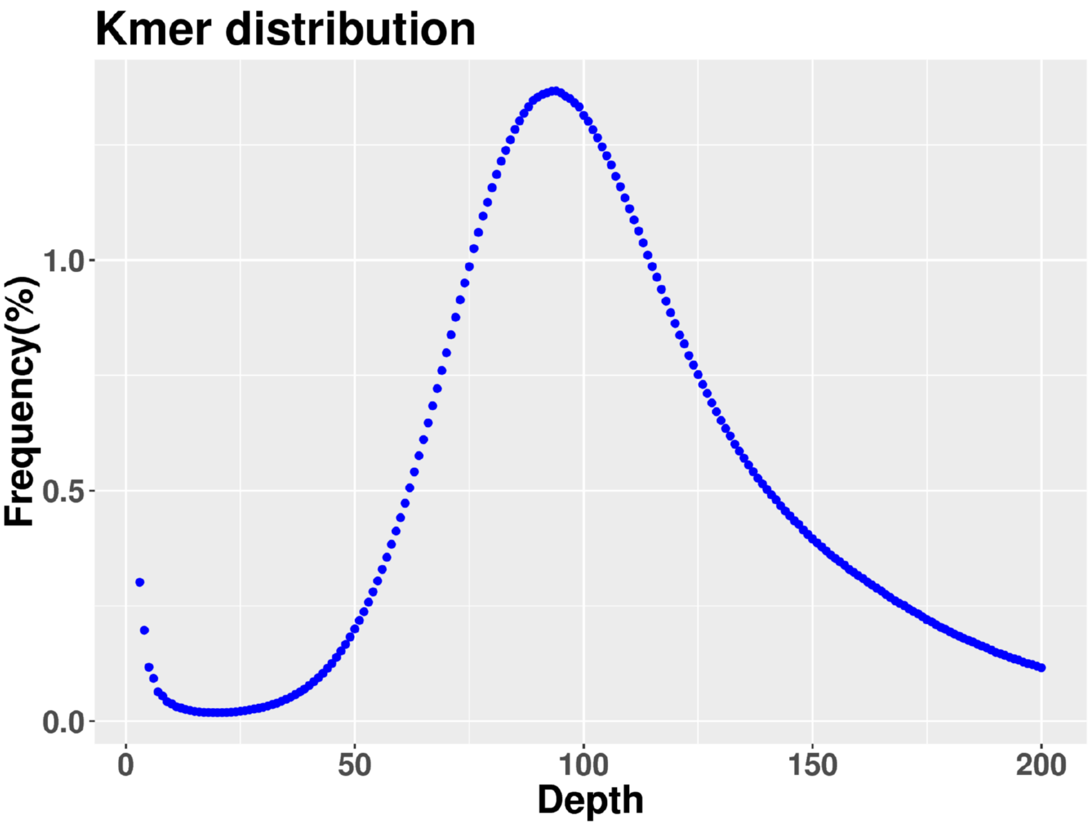


Fig. S1. Genome survey of *P. corylifolia* based on the 21-mer frequency distribution. The distribution was used to estimate a genome size of 484.06 Mb, a heterozygosity rate of 0.03%, and a repetitive sequence content of 51.03%. The x-axis shows K-mer depth, and the y-axis shows frequency (%).

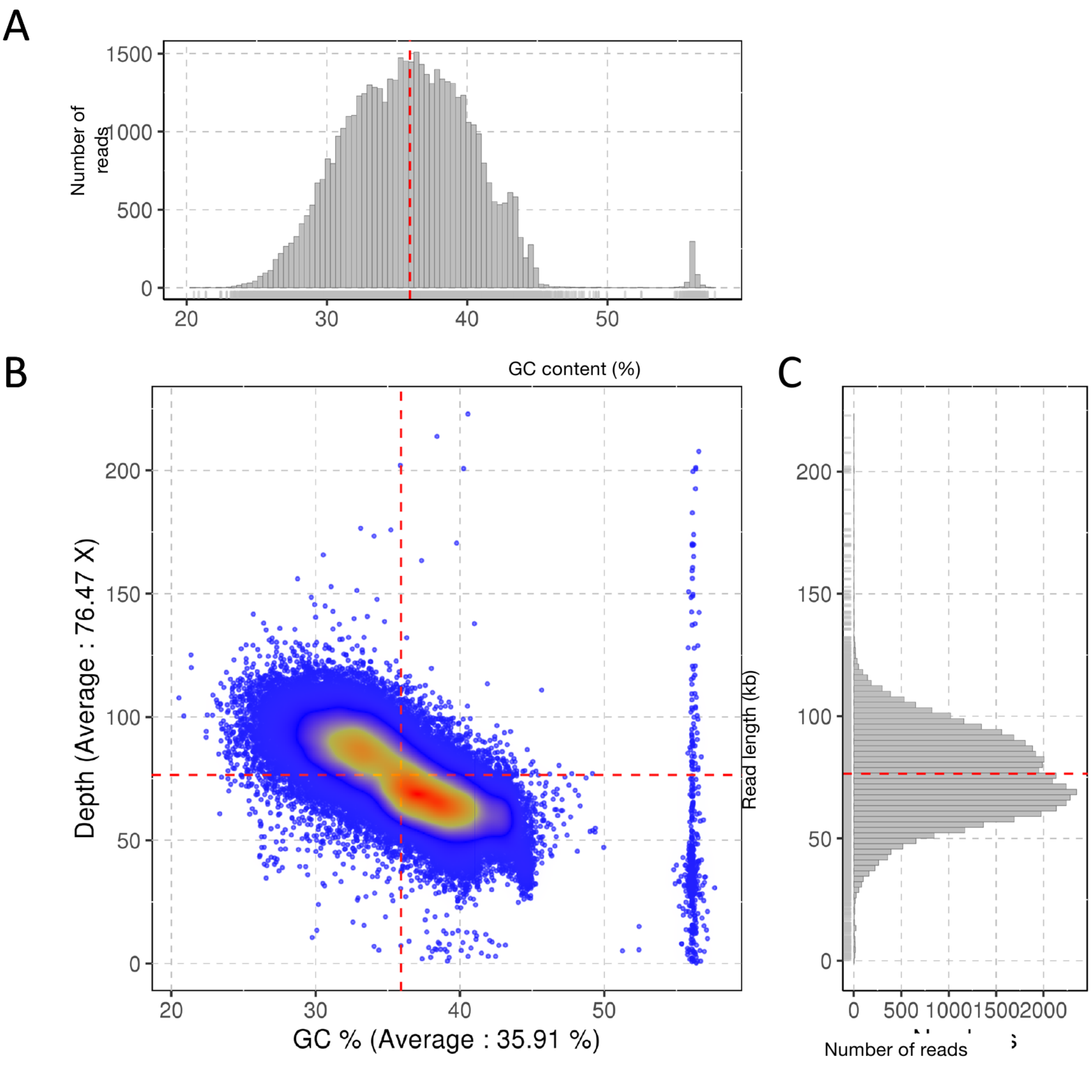


Fig. S2. Quality-control metrics for the PacBio HiFi reads used for genome assembly.

(A) Distribution of GC content (x-axis, %) plotted against the number of reads (y-axis); the average GC content was 35.91%. (B) Relationship between GC content (x-axis, %) and sequencing depth (y-axis, ×); the average sequencing depth was 76.47×. (C) Read-length distribution (y-axis, kb) plotted against the number of reads (x-axis); the mean read length was 14.76 kb and the read N50 was 15.14 kb.

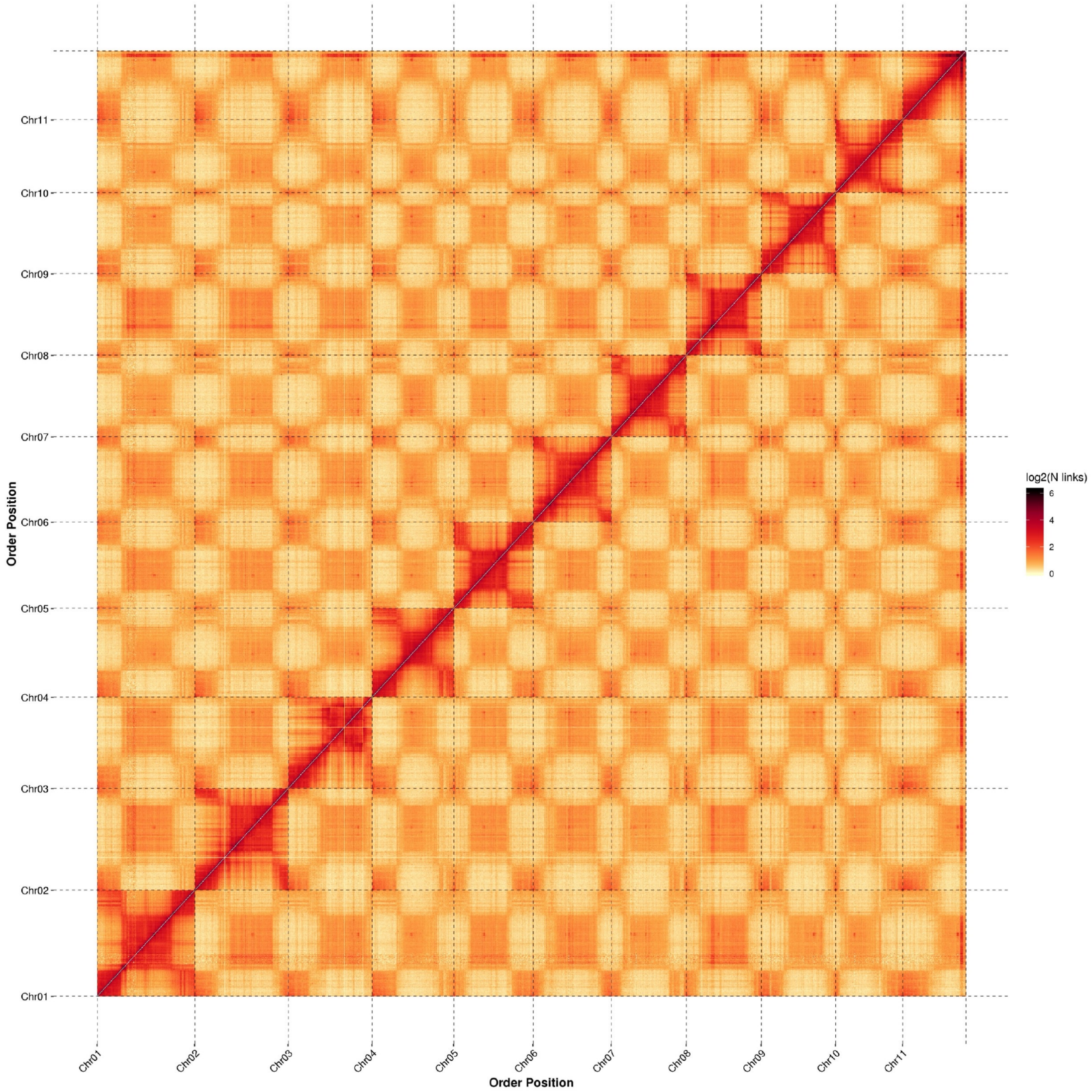


Bins (300kb per bin)

Fig. S3. Hi-C contact map of *P. corylifolia* chromosomes. Pairwise interaction frequencies are shown at 300-kb resolution. The color scale represents log2(N links), with warmer colors indicating higher interaction frequencies. The map was used to anchor contigs to the 11 pseudochromosomes.

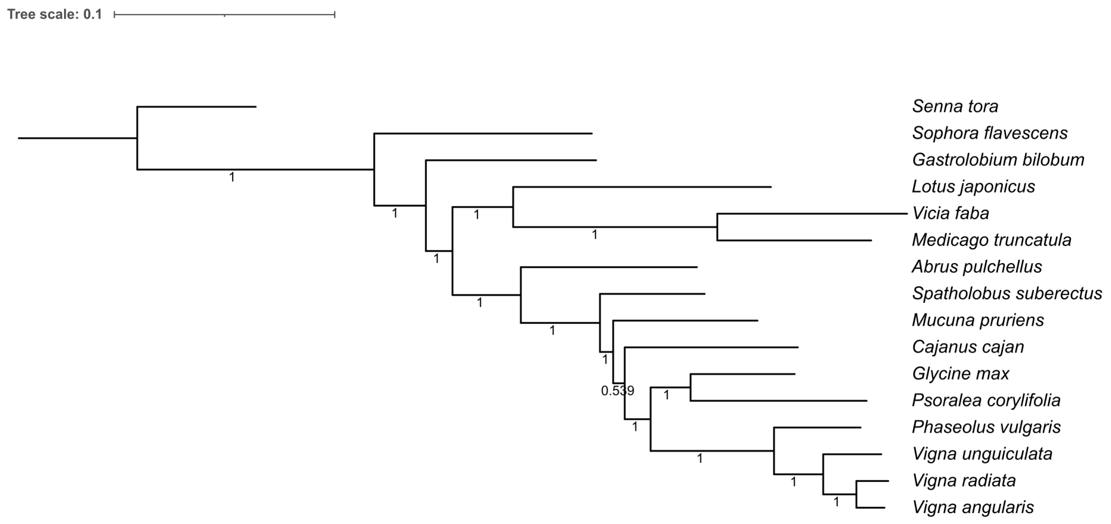


Fig. S4. Phylogenetic tree of *P. corylifolia* and 15 related legume species. The tree was constructed from 117 single-copy orthologous genes using OrthoFinder2/STAG, with *Senna tora* as the outgroup. The scale bar represents 0.1 substitutions per site. The topology places *Glycine max* as the closest relative of *P. corylifolia*.

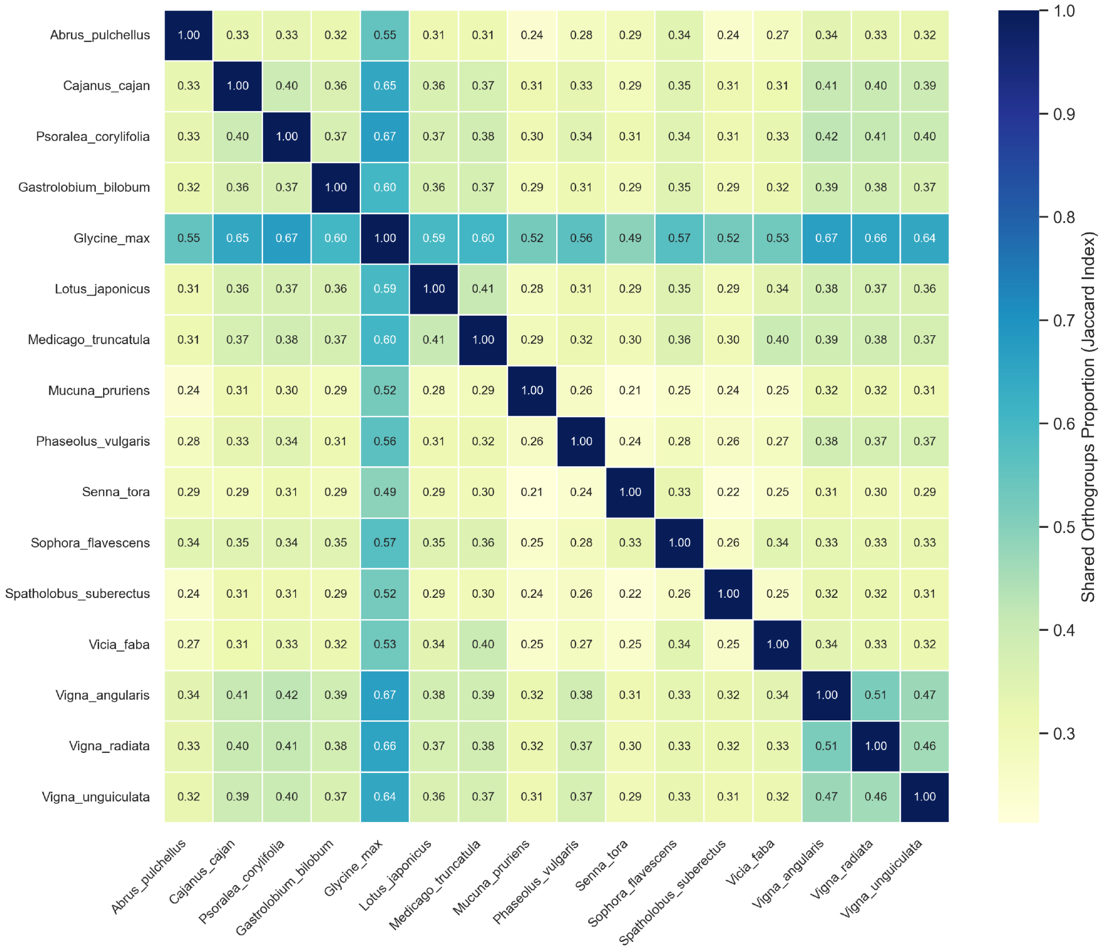


Fig. S5. Species orthogroup overlap heatmap. The heatmap shows the Jaccard similarity of shared orthogroups among 16 legume species. Diagonal values are 1.0, and warmer colors indicate a higher proportion of shared orthogroups. This analysis summarizes the conservation and divergence of orthologous gene sets across the compared species.

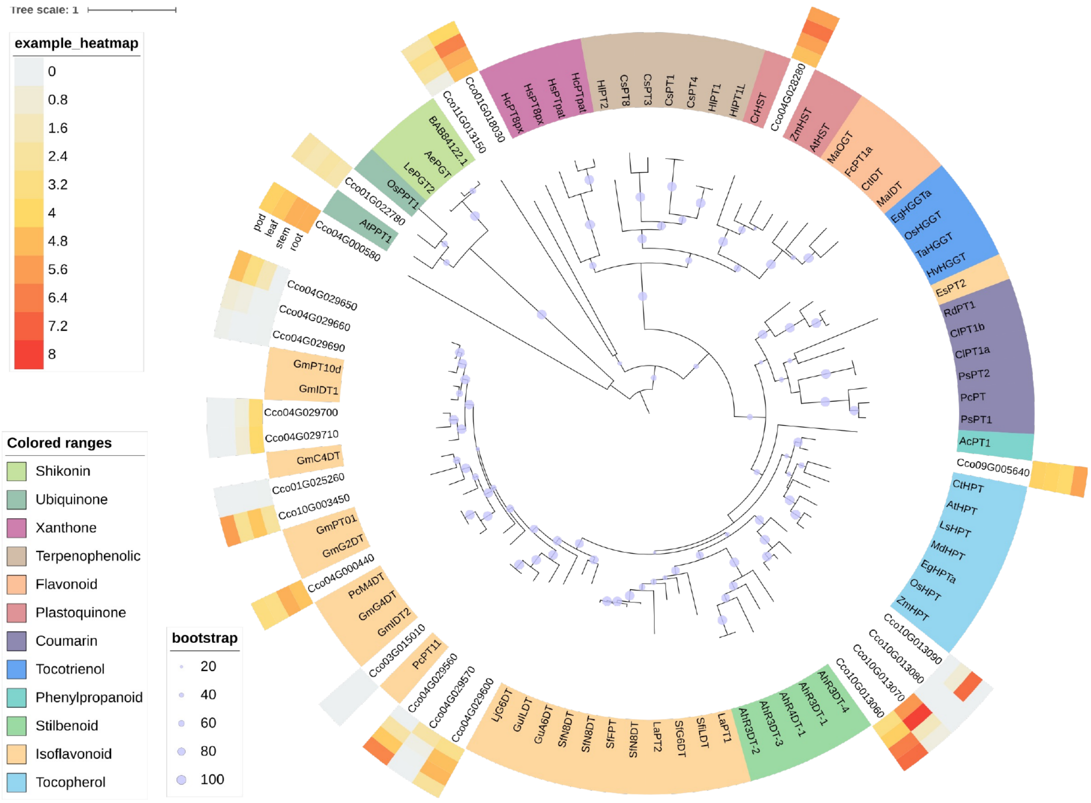


Fig. S6. Phylogenetic relationships and tissue-specific expression of *P. corylifolia* UbiA-type prenyltransferases. The tree includes 22 high-confidence *P. corylifolia* PT candidates and 67 previously characterized plant PTs. Sequences are colored by characterized substrate group, including isoflavonoid, stilbenoid, tocopherol, tocotrienol, xanthone, shikonin, coumarin, flavonoid, plastoquinone, ubiquinone, terpenophenolic, and phenylpropanoid groups. Blue points indicate bootstrap support values (20-100). The adjacent heatmap summarizes FPKM expression across *P. corylifolia* tissues, with the expression scale shown on the figure. The tree scale bar represents 1 substitution per site. FPKM expression was measured in n = 3 biologically independent samples per tissue.

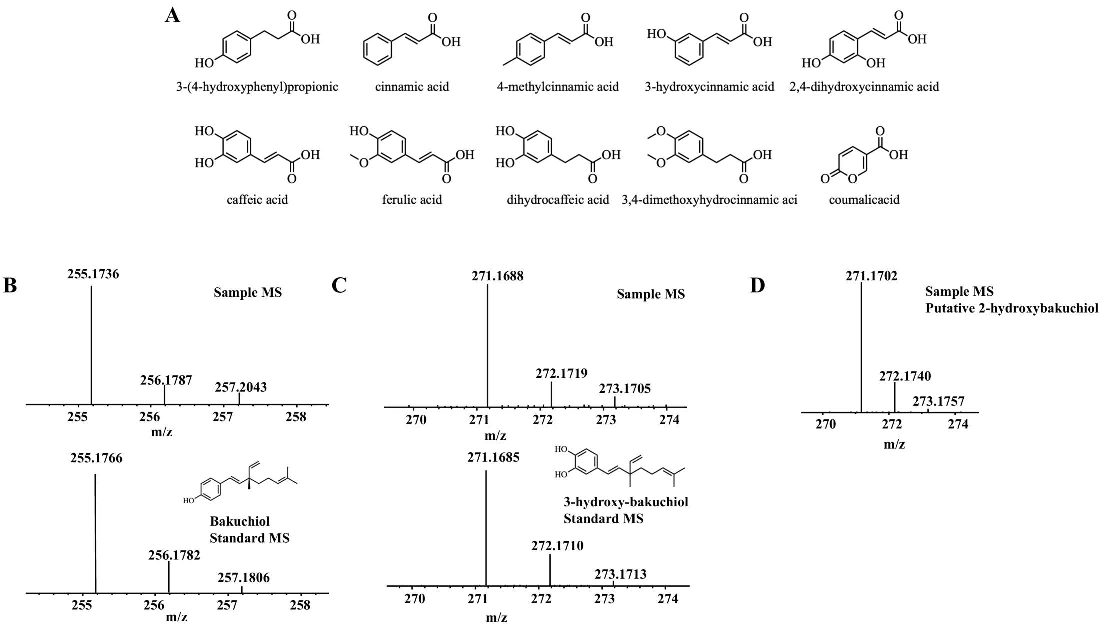


Fig. S7. Substrate specificity of PcPT6. (A) Chemical structures of the substrate analogues tested to evaluate PcPT6 substrate specificity. (B) Mass spectra of the BAK product (Sample MS) and an authentic BAK standard (Standard MS). (C) Mass spectra of the product obtained after feeding caffeic acid (Sample MS) and an authentic 3-hydroxybakuchiol standard (Standard MS). (D) Mass spectrum of the new product obtained after feeding 2,4-dihydroxycinnamic acid. This product was putatively assigned as 2-hydroxybakuchiol based on its mass spectrum, the substrate structure, and the catalytic characteristics of PcPT6. An authentic 2-hydroxybakuchiol standard was unavailable; therefore, the structural assignment in (D) remains tentative.

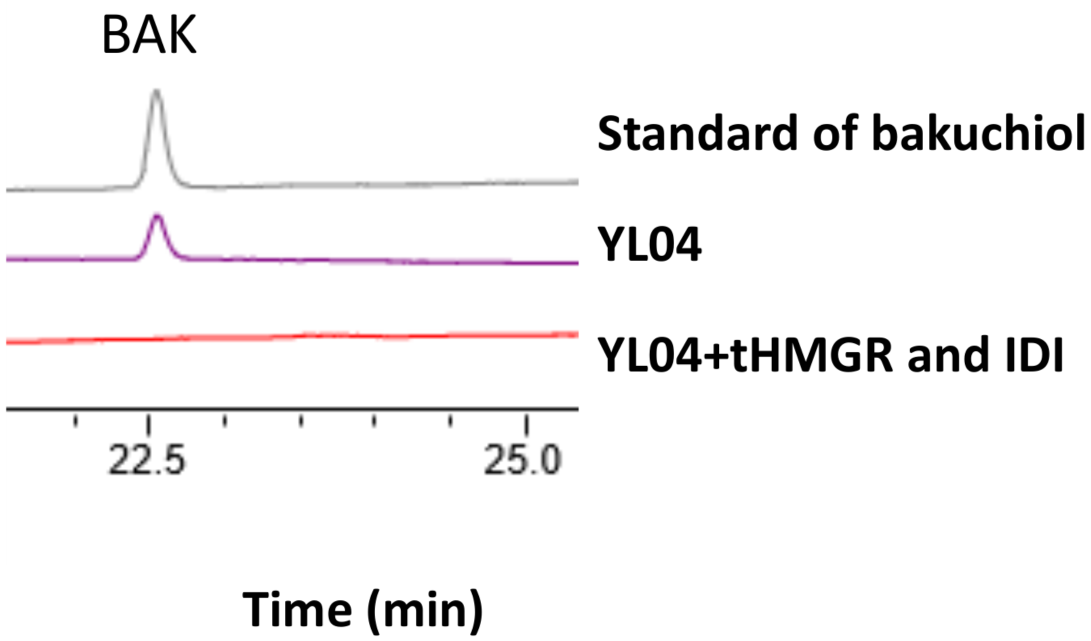


Fig. S8. HPLC analysis of BAK production in strain YL04 and YL04 overexpressing *tHMGR* and *IDI*. An authentic BAK standard is shown for comparison. No BAK peak was detected in the *tHMGR*/*IDI*-overexpressing strain under the tested conditions, indicating that reinforcing this part of the MVA pathway alone was insufficient in the tested background.

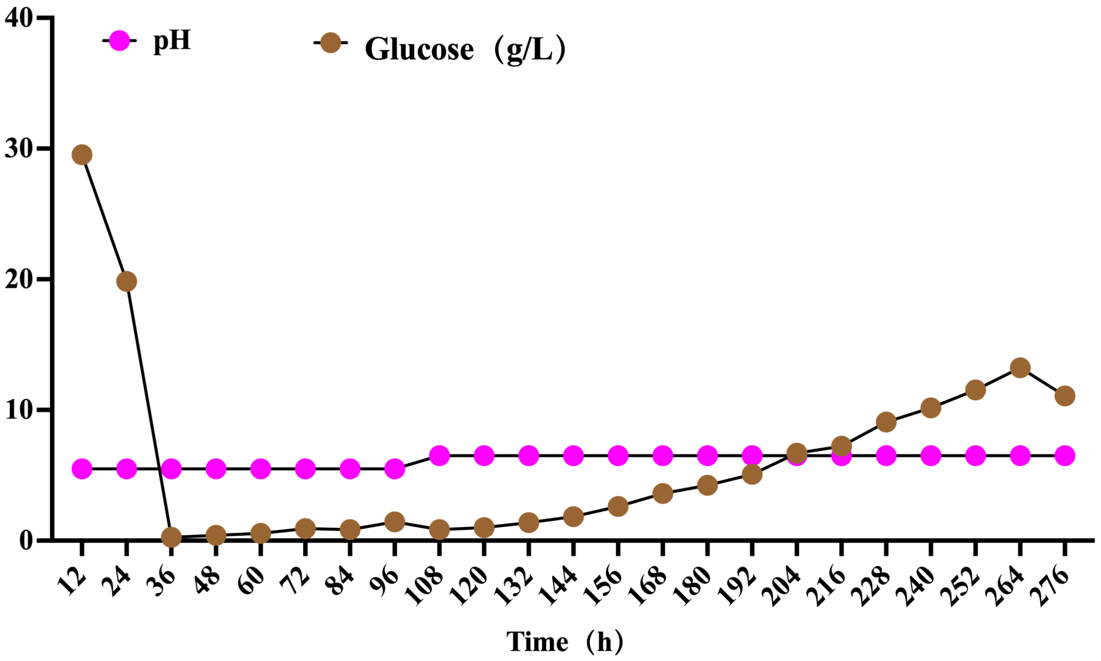


Fig. S9. Profiles of pH and residual glucose concentration during 7.5-L fed-batch fermentation of strain YL23. Samples were analyzed over time; the pH was adjusted to 6.5 after 96 h, and residual glucose reached 13 g/L at 252 h. These profiles indicate that glucose depletion was not the main limitation during the later stage of fermentation.

**Primers used in this study**

| Name | Prime |
|---|---|
| t30PcPT6-F | actttttacaacaaatataaaaacaatgtaccacgctacctcctaca |
| t40PcPT6-F | caaatataaaaacaatgtcctggaacaagactgaaaaga |
| t45PcPT6-F | actttttacaacaaatataaaaacaatggaaaagatccaacacgaatacgc |
| t60PcPT6-F | actttttacaacaaatataaaaacaatgaacttgaagcacaactacggt |
| t70PcPT6-F | actttttacaacaaatataaaaacaatgggtgtctcgacttgtgaaga |
| t90PcPT6-F | actttttacaacaaatataaaaacaatgtccttggaatcccacgaa |
| t110PcPT6-F | actttttacaacaaatataaaaacaatgagattctgtgatgctttctacag |
| PcPT6-R | tgacataactaattacatgatcaacggattagagggatcatca |
| CYC-F | tcatgtaattagttatgtcacgcttaca |
| PGK pro-R | tgtttttatatttgttgtaaaaagtagataattacttccttga |
| tVrPT1-F | caaggaagtaattatctactttttacaacaaatataaaacaatggcttggtgtaagactgaaaaaat |
| tVrPT1-R | gtaagcgtgacataactaattacatgatcatctcatgaatgggataatcaaatattcag |
| tVrPT2-F | caaggaagtaattatctactttttacaacaaatataaaacaatgtccagatacaagaccgaaaaga |
| tVrPT2-R | gtaagcgtgacataactaattacatgatcaacgtaacagcaagatgataatgtattc |
| tVrPT3-F | caaggaagtaattatctactttttacaacaaatataaaacaatgacccgttacaagaccga |
| tVrPT3-R | gtaagcgtgacataactaattacatgatcaaatgatgaatgggataatcaagtattcagc |
| Y33-TDH3-5F | tttgtttgtttatgtgtgtttattcga |
| Y33-PDC1-3R | gcgatttaatctctaattattagttaaagt |
| ScERG20-F(Gibson) | agtttcgaataaacacacataaacaaacaaaatggcttcagaaaaagaaattaggag |
| ScERG20-R(Gibson) | actttaactaataattagagattaaatcgcctatttgcttctcttgtaaactttgttc |
| Sc-Amp-Gibson-F | acccagaaaacgctggtgaaag |
| Sc-Amp-Gibson-R | cgaccaaccagtaagcctgc |
| F96W-F | gcaggcttactggttggtcg |
| F96W-R | ctttcaccagcgtttctgggt |
| N127W-F | aatggccatctgggacgcatt |
| N127W-R | aatggctcccagatggcaatt |
| CluTAL-F | ttttgcagtactaaccgcagatgaagatcaacaacttcct |
| CluTAL-R | aggttgattccgaacagaagttacatggccttcaggtact |
| RgPAL-F | ttttgcagtactaaccgcagatggccccttccgtggactc |
| RgPAL-R | gaggttgattccgaacagaagttaggccatcatcttgaccag |
| EbC4H-F | ctttttgcagtactaaccgcagatggacctgctgttgctcga |
| EbC4H-R | cttgaggttgattccgaacagaagtcagaaggaccttggcttagcg |
| ATR2-F | acgtggggacaggccatggatcaccacacgtctcggagat |
| ATR2-R | tgaattaaacacacatcaacatgtcctcaagctcgtcgag |
| YlCYB5-F | ttttgcagtactaaccgcagatgatcatcaacggcaaggt |
| YlCYB5-R | aggttgattccgaacagaagttatttctgaccctggaggt |

| | |
|---|---|
| ARO4-F | cacacatacaaccacacacatccacatgtcccgttcctcctctccca |
| ARO4-R | gatccttagtttcgggttcccacttagttcttgtttcgtcgctcct |
| ARO7-F | cacacatacaaccacacacatccacatggacttcactaaagccgaca |
| ARO7-R | ggatccttagtttcgggttcccacctactccaaccgccggagcag |
| $ARO4^{K221L}$-F | gatggcggcaacaccctgcagggtgacacccatgaagtg |
| $ARO4^{K221L}$-R | cacttcatgggtgtcaccctgcagggtgttgccgccatc |
| $ARO7^{G139S}$-F | gagatcagcccgagaactattcgtcggtcatggtgtgcg |
| $ARO7^{G139S}$-R | cgcacaccatgaccgacgaatagttctcgggctgatctc |
| ARO1-F | cactttttgcagtactaaccgcagatgtttgccgagggtcagat |
| ARO1-R | ccttgaggttgattccgaacagaagttagttaccaagaacagccttctctcc |
| ARO2-F | ctcacttccccatccacacttttagaaaaaggactttgcgccc |
| ARO2-R | cttgaattaaacacacatcaacatgagcactttcggcacg |
| ARO3-F | actttttgcagtactaaccgcagatgcccgctatgcacaacg |
| ARO3-R | ccttgaggttgattccgaacagaagctaacctcgtcgagtcttgacgg |
| $ARO3^{K225L}$-F | ccacttcctttccgtcaccctgcccggtg |
| $ARO3^{K225L}$-R | gtgacggaaaggaagtggtgg |
| $AroG^{*}$-F | gcagtactaaccgcagatgaactaccagaacgatgatctgc |
| $AroG^{*}$-R | gaggttgattccgaacagaagtcagccacgccgag |
| $TyrA^{*}$-F | cttccccatccacactttcattgccggttatcgttgg |
| $TyrA^{*}$-R | cttgaattaaacacacatcaacatggtagccgagctgacg |
| tHMGR-F | accagcactttttgcagtactaaccgcagatgacccagtctgtgaaggtggttg |
| tHMGR-R | cgttaaccttgaggttgattccgaacagaagctatgaccgtatgcaaatattc |
| IDI-F | cttccccatccacacttctacttgatccaccgccgaatc |
| IDI-R | ttcattcttgaattaaacacacatcaacatgacgacgtcttacagcg |
| MAF1-F | cttccccatccacacttctactccagatcaagttctccg |
| MAF1-R | cttgaattaaacacacatcaacatgcagttcatacaagatcacacac |
| t40PcPT6-F | gcagtactaaccgcagatggcttcttggaacaagaccgaga |
| t40PcPT6-R | cttgaggttgattccgaacagaagtcaacggatgagtggaatcatc |
| $GgaFPS^{N144W}$-F | gcagtactaaccgcagatgcacaagttcactggagtca |
| $GgaFPS^{N144W}$-R | tgattccgaacagaagtttttgacgcttgtagatcttctgg |
| ERG13-F | ctttttgcagtactaaccgcagatgtcgcaaccccagaacgttg |
| ERG13-R | ttgaggttgattccgaacagaagctactgcttgatctcgtac |
| ERG10-F | cttgaattaaacacacatcaacatgcgactcactctgccccg |
| ERG10-R | gcactcacttccccatccacacttctactcgacagaagagaccttcttg |
| ERG19-F | ctttttgcagtactaaccgcagatgatccaccaggcctccaccac |
| ERG19-R | cttgaggttgattccgaacagaagctacttgctgttcttcagagaac |
| ERG8-F | ctttttgcagtactaaccgcagatgaccacctattcggctcc |
| ERG8-R | aggttgattccgaacagaagctacttgaaccccttctcgagc |
| ERG12-F | tcacttccccatccacacttctaatgggtccagggaccg |
| ERG12-R | tgaattaaacacacatcaacatggactacatcatttcggcgc |
| YL-P1-F | tacttactcaagtacaaataaatatcgatactaatacgag |

| | |
|---|---|
| YL-P1-R | actatgagaacgtgtctagcgaag |
| YL-P2-F | aattcgacgacctgaacactcg |
| YL-P2-R | gattgaaacgcctgacaaaaacgc |
| YL-P3-F | aaaaaactgtagtagtgtggtgatgga |
| YL-P3-R | aattattgcacaggacacacacaag |
| YL-P5-F | ttgatatggtgtaacaatgataaaccaaggc |
| YL-P5-R | gttagaagcaattggagaagaaacgttcag |
| YL-P6-F | ttgatatggtgtaacaatgataaaccaaggc |
| YL-P6-R | gttagaagcaattggagaagaaacgttcag |
| YL-P7-F | gcggaggagcaatagacatacga |
| YL-P7-R | ttaacactggaccgtactgccca |
| YL-P8-F | ataatggatgcctaaggagcccggaaga |
| YL-P8-R | gtatgtacgatagtcgtctgatcaggtgggag |
| YL-P9-F | atgttttaaacagtgtggcctcattactttgatcg |
| YL-P9-R | gacctgaggttgttgatgcgccatt |
| YL-P12-F | agatagagtctcttcaactcgctcaaaaaacaacc |
| YL-P12-R | atcgtctggcagatggccttggacg |
| YL-P13-F | tggtgattacttgaatatagttgaacaccattc |
| YL-P13-R | tgcttgtacgtctcaaatttttttattttttttatgg |
| YL-P15-F | ttggccgattgccgctcagaggcatta |
| YL-P15-R | tcttgatgcggaatcggtctagagtcctagctgatg |
| YL-P16-F | tattggtcgttgtcagaccatcgagtcga |
| YL-P16-R | aggtagggtgggggaagagtgaaca |
| YL-P18-F | aggcaagtcgctgtctgtgttcgat |
| YL-P18-R | gcctgtaagctaatatgaaggggggtgggg |
| YL-P19-F | atgctgcgggcgg |
| YL-P19-R | tctgatggcctggagcgag |
| YL-P20-F | tgggagtgtatttggaaaatgact |
| YL-P20-R | tcttgtaattccatagataatatatacgaaacgca |
| YL-P22-F | gccgacaatctctcttcctc |
| YL-P22-R | taaaacaatgacaaatacaactctagactcca |
| YL-pMHY1-F | tattatgtcgtcgaaaaaggaagtcg |
| YL-pMHY1-R | gcgacgggaggggtctaca |

**Plasmids used in this study**

| Name | Relevant characteristics | Source |
|---|---|---|
| YCplac22-PE | Centromeric vector, *TRP1*, AmpR, pPGK1 and tCYC1 | Genewiz |
| YCplac33-PE | Centromeric vector, *URA3*, AmpR, pTDH3 and tPDC1 | Genewiz |
| Y33-ScERG20$^{F96W, N127W}$ | YCplac33 vector, pTDH3-*ScERG20* $^{F96W, N127W}$-tPDC1 | This study |

| | | |
|---|---|---|
| Y22-PcPT1 | YCplac22 vector, pPGK1-*PcPT1*- tCYC1 | This study |
| Y22-PcPT2 | YCplac22 vector, pPGK1-*PcPT2*- tCYC1 | This study |
| Y22-PcPT3 | YCplac22 vector, pPGK1-*PcPT3*- tCYC1 | This study |
| Y22-PcPT4 | YCplac22 vector, pPGK1-*PcPT4*- tCYC1 | This study |
| Y22-PcPT5 | YCplac22 vector, pPGK1-*PcPT5*- tCYC1 | This study |
| Y22-PcPT6 | YCplac22 vector, pPGK1-*PcPT6*- tCYC1 | This study |
| Y22-t30PcPT6 | YCplac22 vector, pPGK1-*t30PcPT6*- tCYC1 | This study |
| Y22-t40PcPT6 | YCplac22 vector, pPGK1-*t40PcPT6*- tCYC1 | This study |
| Y22-t45PcPT6 | YCplac22 vector, pPGK1-*t45PcPT6*- tCYC1 | This study |
| Y22-t60PcPT6 | YCplac22 vector, pPGK1-*t60PcPT6*- tCYC1 | This study |
| Y22-t70PcPT6 | YCplac22 vector, pPGK1-*t70PcPT6*- tCYC1 | This study |
| Y22-t90PcPT6 | YCplac22 vector, pPGK1-*t90PcPT6*- tCYC1 | This study |
| Y22-t110PcPT6 | YCplac22 vector, pPGK1-*t110PcPT6*- tCYC1 | This study |
| Y22-tVrPT1 | YCplac22 vector, pPGK1-*tVrPT1*- tCYC1 | This study |
| Y22-tVrPT2 | YCplac22 vector, pPGK1- *tVrPT2*- tCYC1 | This study |
| Y22-tVrPT3 | YCplac22 vector, pPGK1- *tVrPT3*- tCYC1 | This study |
| P1-(tLIP2-tPEX20) | PMD-19T, AmpR, tLIP2, tPEX20 Homologous arms *KU80*-UP and *KU80*-DN | Wang et al. |
| P2-(tLIP2-tPEX20) | PMD-19T, AmpR, tLIP2, tPEX20 Homologous arms IntC2-UP and IntC2-DN | Wang et al. |
| P3-(tLIP2-tPEX20) | PMD-19T, AmpR, tLIP2, tPEX20 Homologous arms IntC3-UP and IntC3-DN | Wang et al. |
| P5-(tLIP2-tPEX20) | PMD-19T, AmpR, tLIP2, tPEX20 Homologous arms IntE1-UP and IntD1-DN | Wang et al. |
| P6-(tLIP2-tPEX20) | PMD-19T, AmpR, tLIP2, tPEX20 Homologous arms IntE3-UP and IntE3-DN | Wang et al. |
| P7-(tLIP2-tPEX20) | PMD-19T, AmpR, tLIP2, tPEX20 Homologous arms IntE4-UP and IntE4-DN | Wang et al. |
| P8-(tLIP2-tPEX20) | PMD-19T, AmpR, tLIP2, tPEX20 Homologous arms *ARO10*-UP and *ARO10*-DN | Wang et al. |
| P9-(tLIP2-tPEX20) | PMD-19T, AmpR, tLIP2, tPEX20 Homologous arms *PDC5*-UP and *PDC5*-DN | Wang et al. |
| P12-(tLIP2-tPEX20) | PMD-19T, AmpR, tLIP2, tPEX20 Homologous arms D17-UP and D17-DN | Wang et al. |
| P13-(tLIP2-tPEX20) | PMD-19T, AmpR, tLIP2, tPEX20 Homologous arms B3-UP and B3-DN | Zhu et al. |
| P15-(tLIP2-tPEX20) | PMD-19T, AmpR, tLIP2, tPEX20 Homologous arms E14-UP and I E14-DN | Zhu et al. |
| P16-(tLIP2-tPEX20) | PMD-19T, AmpR, tLIP2, tPEX20 Homologous arms F17-UP and F17-DN | Zhu et al. |
| P19-(tLIP2-tPEX20) | PMD-19T, AmpR, tLIP2, tPEX20 Homologous arms *DGA1*-UP and *DGA1*-DN | Zhu et al. |
| P20-(tLIP2-tPEX20) | PMD-19T, AmpR, tLIP2, tPEX20 Homologous arms *DGA2*-UP and *DGA2*-DN | Zhu et al. |
| P22-(tLIP2-tPEX20) | PMD-19T, AmpR, tLIP2, tPEX20 Homologous arms *ARE1*-UP and *ARE1*-DN | This study |
| PMHY1-(tLIP2-tPEX20) | PMD-19T, AmpR, tLIP2, tPEX20 Homologous arms *MHY1*-UP and *MHY1*-DN | This study |

| | | |
|---|---|---|
| G1 | pNat-YLgRNA_*KU80* | Wang et al. |
| G2 | pNat-YLgRNA_IntC2 | Wang et al. |
| G3 | pNat-YLgRNA_IntC3 | Wang et al. |
| G5 | pNat-YLgRNA_IntD1 | Wang et al. |
| G6 | pNat-YLgRNA_IntE3 | Wang et al. |
| G7 | pNat-YLgRNA_IntE4 | Wang et al. |
| G8 | pNat-YLgRNA_*ARO10* | Wang et al. |
| G9 | pNat-YLgRNA_*PDC5* | Wang et al. |
| G12 | pNat-YLgRNA_D17 | Wang et al. |
| G13 | pNat-YLgRNA_B3 | Zhu et al. |
| G15 | pNat-YLgRNA_E14 | Zhu et al. |
| G16 | pNat-YLgRNA_F17 | Zhu et al. |
| G19 | pNat-YLgRNA_*DGA1* | Zhu et al. |
| G20 | pNat-YLgRNA_*DGA2* | Zhu et al. |
| G22 | pNat-YLgRNA_*ARE1* | This study |
| G-MFE1 | pNat-YLgRNA_*MFE1* | Zhu et al. |
| G-MHY1 | pNat-YLgRNA_*MHY1* | This study |
| P18-CluTAL-RgPAL | P18 vector, pTEFIN-*CluTAL*-tLIP2/pGPD-*RgPAL*-tPEX20 | This study |
| P5-YlCYB5-ATR2-EbC4H | P5 vector, pTEFIN-*YlCYB5*-tLIP2/pGPD-*ATR2*-tXPR2/pGPD-*EbC4H*-tPEX20 | This study |
| P15-tHMGR-IDI | P15 vector, pTEFIN-*tHMGR*-tLIP2/pGPD-*IDI*-tPEX20 | This study |
| P3-PcPT6 | P3 vector, pTEFIN-*PcPT6*-tLIP2 | This study |
| P7-ARO4$^{K221L}$-ARO7$^{G139S}$ | P7 vector, pTEFIN- *ARO4*$^{K221L}$ -tLIP2/pGPD- *ARO7*$^{G139S}$ -tPEX20 | This study |
| P6-ARO1-ARO2 | P6 vector, pTEFIN- *ARO1* -tLIP2/pGPD-*ARO2*-tPEX20 | This study |
| P1-ARO3$^{K225L}$ | P1 vector, pTEFIN- *ARO3*$^{K225L}$ -tLIP2 | This study |
| P2-AroG$^{*}$-TyrA$^{*}$ | P2 vector, pTEFIN- *AROG*$^{*}$ -tLIP2/pGPD-*TyrA*$^{*}$-tPEX20 | This study |
| pMHY1-tHMGR-IDI | PMHY1 vector, pTEFIN- *tHMGR* -tLIP2/pGPD- *IDI*-tPEX20 | This study |
| P2-MAF1 | P2 vector, pTEFIN-*MAF1*-tLIP2 | This study |
| P8-t40PcPT6 | P8 vector, pTEFIN- *t40PcPT6*-tLIP2 | This study |
| P9-t40PcPT6 | P9 vector, pTEFIN- *t40PcPT6*-tLIP2 | This study |
| P22-AgrGPPS2 | P22 vector, pTEFIN- *AgrGPPS2*-tLIP2 | This study |
| P22-ScERG20$^{WW}$ | P22 vector, pTEFIN- *ScERG20*$^{WW}$-tLIP2 | This study |
| P22-GgaFPS$^{N144W}$ | P22 vector, pTEFIN- *GgaFPS*$^{N144W}$ -tLIP2 | This study |
| P19-t40PcPT6 | P19 vector, pTEFIN- *t40PcPT6* -tLIP2 | This study |
| P12-ERG10-ERG13 | P12 vector, pTEFIN- *ERG10* -tLIP2/pGPD-*ERG13*-tPEX20 | This study |
| P13-ERG19 | P13 vector, pTEFIN-*ERG19*-tLIP2 | This study |
| P16-ERG8-ERG12 | P16 vector, pTEFIN- *ERG8* -tLIP2/pGPD-*ERG12*-tPEX20 | This study |
| P20-t40PcPT6 | P20 vector, pTEFIN- *t40PcPT6* -tLIP2 | This study |

**Strains used in this study**

| Strain | Genotype | Reference |
|---|---|---|
| W303-1B | MATα, *ade2–1; ura3–1; his3–11,15; trp1–1;* | Stored in lab |

| | | |
|---|---|---|
| | *leu2–3,112; can 1–100* | |
| *E.coli* DMT | F$^-$; φ80 *lac* Z*Δ*M15 *Δ*(*lac*ZYA-*argF*) U169 *rec*A1 *end*A1 *hsd*R17($r_K^-$,$m_K^+$) *phoA sup*E44 thi-1 *gyr*A96 *rel*A1 *tonA* | TransGen |
| ScP00 | W303 carrying YCplac33 vector and YCplac22 vector | This study |
| ScP01 | W303 carrying Y33-*ScERG20*$^{F96W,\ N127W}$ and Y22-*PcPT1* | This study |
| ScP02 | W303 carrying Y33-*ScERG20*$^{F96W,\ N127W}$ and Y22-*PcPT2* | This study |
| ScP03 | W303 carrying Y33-*ScERG20*$^{F96W,\ N127W}$ and Y22-*PcPT3* | This study |
| ScP04 | W303 carrying Y33-*ScERG20*$^{F96W,\ N127W}$ and Y22-*PcPT4* | This study |
| ScP05 | W303 carrying Y33-*ScERG20*$^{F96W,\ N127W}$ and Y22-*PcPT5* | This study |
| ScP06 | W303 carrying Y33-*ScERG20*$^{F96W,\ N127W}$ and Y22-*PcPT6* | This study |
| ScP07 | W303 carrying Y33-*ScERG20*$^{F96W,\ N127W}$ and Y22-*t30PcPT6* | This study |
| ScP08 | W303 carrying Y33-*ScERG20*$^{F96W,\ N127W}$ and Y22-*t40PcPT6* | This study |
| ScP09 | W303 carrying Y33-*ScERG20*$^{F96W,\ N127W}$ and Y22-*t45PcPT6* | This study |
| ScP10 | W303 carrying Y33-*ScERG20*$^{F96W,\ N127W}$ and Y22-*t60PcPT6* | This study |
| ScP11 | W303 carrying Y33-*ScERG20*$^{F96W,\ N127W}$ and Y22-*t70PcPT6* | This study |
| ScP12 | W303 carrying Y33-*ScERG20*$^{F96W,\ N127W}$ and Y22-*t90PcPT6* | This study |
| ScP13 | W303 carrying Y33-*ScERG20*$^{F96W,\ N127W}$ and Y22-110PcPT6 | This study |
| ScP14 | W303 carrying Y33-*ScERG20*$^{F96W,\ N127W}$ and Y22- *tVrPT1* | This study |
| ScP15 | W303 carrying Y33-*ScERG20*$^{F96W,\ N127W}$ and Y22- *tVrPT2* | This study |
| ScP16 | W303 carrying Y33-*ScERG20*$^{F96W,\ N127W}$ and Y22- *tVrPT3* | This study |
| YL00 | W29(MATa *KU70*Δ:: PrTEF1-cas9-TTef12:: PrGPD-hyg-TLip2) | Stored in lab |
| YL01 | YL00 E13:: pTEFIN-*CluTAL*-tLIP2/pGPD-*RgPAL*-tPEX20 | This study |
| YL02 | YLP01 IntE1:: pTEFIN-*YlCYB5*-tLIP2/pGPD-*ATR2*-tXPR2/pGPD-*EbC4H*-tPEX20 | This study |
| YL03 | YL02 E14:: pTEFIN-*tHMGR*-tLIP2/pGPD-*IDI*-tPEX20 | This study |
| YL04 | YL03 IntC3:: pTEFIN-*PcPT6*-tLIP2 | This study |
| YL05 | YL04 Δ*MFE1* | This study |
| YL06 | YL05 IntE4:: pTEFIN- *ARO4*$^{K221L}$ -tLIP2/pGPD- *ARO7*$^{G139S}$ -tPEX20 | This study |

| | | |
|---|---|---|
| YL07 | YL06 InE3:: vector, pTEFIN- *ARO1* -tLIP2/pGPD- *ARO2*-tPEX20 | This study |
| YL08 | YL07 *KU80*:: pTEFIN- *ARO3*$^{K225L}$ -tLIP2 | This study |
| YL09 | YL08 IntC2:: pTEFIN- *AROG*$^{*}$ -tLIP2/pGPD-*TyrA*$^{*}$-tPEX20 | This study |
| YL10 | YL09 *MHY1*:: pTEFIN- *tHMGR* -tLIP2/pGPD- *IDI*-tPEX20 | This study |
| YL11 | YL09 IntC2:: pTEFIN-*MAF1*-tLIP2 | This study |
| YL12 | YL10 *ARO10*:: pTEFIN- *t40PcPT6*-tLIP2 | This study |
| YL13 | YL12 *PDC5*:: pTEFIN- *t40PcPT6*-tLIP2 | This study |
| YL14 | YL13 *ARE1*:: pTEFIN- *AgrGPPS2*-tLIP2 | This study |
| YL15 | YL13 *ARE1*:: pTEFIN- *ScERG20*$^{WW}$-tLIP2 | This study |
| YL16 | YL13*ARE1*:: pTEFIN- *GgaFPS*$^{N144W}$ -tLIP2 | This study |
| YL17 | YL13 *DGA1*:: pTEFIN- *t40PcPT6*-tLIP2 | This study |
| YL18 | YL16 *DGA1*:: pTEFIN- *t40PcPT6*-tLIP2 | This study |
| YL19 | YL18 *DGA2*:: pTEFIN- *t40PcPT6*-tLIP2 | This study |
| YL20 | YL18 D17:: pTEFIN- *ERG10* -tLIP2/pGPD-*ERG13*-tPEX20 | This study |
| YL21 | YL20 B3:: pTEFIN-*ERG19*-tLIP2 | This study |
| YL22 | YL21 F17:: pTEFIN- *ERG8* -tLIP2/pGPD-*ERG12*-tPEX20 | This study |
| YL23 | YL22 *DGA2*:: pTEFIN- *t40PcPT6*-tLIP2 | This study |

**genes used in this study.**

>ScERG20$^{F96W, N127W}$

MASEKEIRRERFLNVFPKLVEELNASLLAYGMPKEACDWYAHSLNYNTPGGK
LNRGLSVVDTYAILSNKTVEQLGQEEYEKVAILGWCIELLQAYWLVADDMMD
KSITRRGQPCWYKVPEVGEIAIWDAFMLEAAIYKLLKSHFRNEKYYIDITELF
HEVTFQTELGQLMDLITAPEDKVDLSKFSLKKHSFIVTFKTAYYSFYLPVALA
MYVAGITDEKDLKQARDVLIPLGEYFQIQDDYLDCFGTPEQIGKIGTDIQDNK
CSWVINKALELASAEQRKTLDENYGKKDSVAEAKCKKIFNDLKIEQLYHEYE
ESIAKDLKAKISQVDESRGFKADVLTAFLNKVYKRSK*

>PcPT1

MELSLSPTSHHVPSTIPTLNSANLFLKATTPQPLFLGFSKHFNSIGLHHHSHRCC
SNPVPKRPQRPSSIRACTGVGAAGSDRPFAEKLIDFKDACWRFLRPHTIRGTTL
GSFALVARALIENSNLIKWSLLFKALSGLFALICGNGYIVGINQIYDISIDKVNKP
YLPIAAGDLSVQSAWFLVIFFAAAGLSIVGLNFGPFIFSLYTLGLFLGTIYSVPPL
RMKRFPVAAFLIIATVRGFLLNFGVYYATRAALGLAFEWSSPVVFITTFVTFFA
LVIAITKDLPDVEGDRKYQISTFATKLGVRNIAFLGSGILLMNYVVSVLAAIYM
PQAFRQWLLIPAHTIFAVSLIYQARILEQANYTKDAISGFYRFIWNLFYAEYAIF
PFI*

>PcPT2

MSCYLVSVSMNLAFATSSTNACSVTRGNVSQAQHSRKNTYYGSSSTSKASQH
KRNFQIDYPFFRLQNPCLNHYYKCIGLGSTYQECYRKYVVEGTSKSHSDFEPD
ASYPINSSDFVKKIFDALYLFITPYGMVTQVLGIISAALLAVEKLSDISPLFFNGV
LQIIVPNFLMGIYLAGLNQIADVEIDKINKPYLPLASGQLSITTGVIVIASCATMS
LLICWILGSWPLIWGNISLFLIWSAYSFNVPLLRWKKNPFLAALCLFVTWTINL

PISSFFHMQTFVFKRPTIFPKSLSFVAAFMSLYSIGIALCKDIPDVEGDTKYGFYS
FAARYGQKQIFWICLYLFETAFGVALLAGATSPYLWVKIITGLGHVLLASVLW
YQAKATNLTNKASIASFYMLVWKLLCAEYLLMPLIR*
>PcPT3
MNLALATSSTNASSLTRGNVSQTQHSSKNTYDASYYVSKASQHKRKLQIECN
LSRSRQHSLNHHHKSIERARPTFQECSRIYIVKAAPAIQSSDSESHVSGLRNFFD
SAKNFFVILYYFCYPYSMIARTLCTISASFFTVEKLSDISPLFFVGLLQVLAVHFL
MDLYINGVNQVFDFEIDKINKPYLPLPSGKLSFTNAIFITVSSAVMSFWASLIIGS
RPLIWSLVLCFLPLTCYSVNVPLLRWKKYPWLTAILFINSWALVFPITFFLHMQT
FVLKRPAIFPRSLIVSVVFLSLYSIGIALSKDIPDIEGDKKFGIHSFSARLGQKQV
FWICVSFFETAFGVAMLAGATSSASIWIKLITGLGNIILASILWYQTQYVDLTSP
ASTRSFYMLIWKLLYAAYFLLPFIR*
>PcPT4
MDSLLLRSFPNNASSSFFATANAADFSRTRTKCFANIYYASSYVPNVSWHNRKI
HKEYNFLRFGWPSLNHHYKGIEGGCTCKKCNTKFVVKATSEKSFESEPQALD
PKSILNSVQNSLDAFYRFSRPHTVIGTALSIISVSLLAVEKISDISPLFFTGVLEAV
VAALFMNIYIVGLNQLSDVEIDKINKPYLPLASGEYSFETGVAIVASFSILSFWL
GWIVGSWPLFWALFVSFVLGTAYSINVPLLRWKRFAVLAAMCILAVRAVIVQL
AFFLHMQTHVYKRPPVFSRPLIFATAFMSFFSVVIALFKDIPDIEGDKVFGIQSF
SVRLGQKPVFWTCVALLEIAYGVALLVGAASPCLWSKIVTGLGHALLASILCY
RAKSVDLKSKASITSFYMFIWKLFYAEYLLIPFVR*
>PcPT5
MNSGFLLSSPNNASSSLTTTGSLTCVNQALNGPVRGLGSIYSPLCSYAAKASQN
RSKTKIKYTAMQLQHPSLNHHNKWIEGESTYQECKKKYVVQAVSESAFDSEP
HASDPKMTYLDSVKDLLAIIYKFSYPYAMIGIILCAVSSSLLTVEKLSDISPSFIV
GVLQAAIPHVMIAIYSNSINQVFDLQIDRINKPHLPLASGQLSTKTVAIIAASFLI
MSFGLSWSIGSWPLIWNVICTFSIWTAYSMNAPLLRWKKNPLLAAFCMVSSW
GFVLPITFYLHMQTFVLKRPSVFPTSLIFVVAFMSLYSLGMALAKDIPDVEGDK
LCGVDTFSIRWGPKQVFWICVLLFEMAFGIALLGGATSPNPWVKLITLVGNTA
LASVLWYQAKSTDLSNPASGQSFYMLNWKLLYAGYILIGLIR*
>PcPT6
MASMFLGSLPLASSANYIGRSTRSKKCTESYHATSYITTASWNKTEKIQHEYA
NMRHRQHNLKHNYGGIEGVSTCEDWARNFVVNAASGESLESHEAQHHTPET
LLGSIKRFCDAFYRFSRPHVITGTAINIIVMSSLALENSSDISTKFFIGLFQAILTTI
PMNLYASGLNQLTDIEIDKINKPYLPLASGEFSVRTGVIIITLCAIMSLLVGWVV
GSPALFWTSFAYFVLGTVYTINLPLLRWKRYPAFAALCFFIIRGLMFHVAFFLHI
QTHVFKRPMMIPKSVMFGTAFMSFFYVIIALFKDIPDIEGDKEHGVKSLTMRL
GQERVFWICVSLLLMAYGGAIVVGATSSFLWSKLITVSGHALLASIFWNRANS
VDLKSHEEITSLYMFMWKLFYIEYLMIPLIR*
>tVrPT1
MAWCKTEKIQNKYHSMRHHQNHHTGIEVGSTFHQGARRYVLNAASGESFES
AKPKHFPKPFRGSIKHYLDAFTRFFNVYVLIGSAISITSMSLLAVEKLSDISLTYF
IGLFQAQVAAALMSGYVIGLNHLTDVEIDKINKPYRPLASGEYSFGTGLILTAS
CLILSFGIGWIVGSPPLLWALSIHFLLGTAYSIDLPLLRWKRIAVLAAMCIVVVH

GVMFQLAYFLHMQTFVLQRPIMLPRSLLFSTAFLSFFSLVLALFKDIPDHEGDE
KHGIKSLSVLLGQKRVFWICVSLLEMIYGVAILVGATSSNHWSKMITVSAHAV
LASILLYHSKSVDVKSSSATASFYMFLWKLFYAEYLIIPFMR*
>tVrPT2
MSRYKTEKIQNKYHIVRRQKHYRGIEVGSCHQWAKSFAVDAAPGDLFESDKP
QHFPKPFWGSVKYYLYTFSRFTKPYVSIGTAISITSMALLTVEKLSDISLTYFIGL
FQAIVAATLMCSYVIGLNNLTDVEIDKINKPNRPLASGEYSFETGVILTAACLIL
SFGIGWIVGSPPLLWALSIHFLLGTAYSIDLPLLRWKRIAALAAMCIVVVQGIIF
QVAYFLHMQTYVLKRPTMLPRSVIFATAFMSFFCLVLALFKDIPDTEGDEKFGI
QSLSVHLGQKRVFWICVSLLEMIYGVAILVGATSSRLWSKMIMVLAHALLASI
LWYHAKSVDLKSNSAIESFYMFIWKLFYAEYIIILLLR*
>tVrPT3
MTRYKTEKIQNNYHIMRHHQNHYSSIKVGSTFHQRARRYVVNAASGESFESA
KPQHFQKTFTGSIKYYLDAFTRFFKPYVLIAAAISITSMSLLAVEKLSDISLTYFI
GLFQAIVAAALMACYITGLNHLTDVEIDKINKPYRPLASGEYSFETAVILTAASL
ILSFGIGWIVGSPPLLWALSITFLLGTAYSIDLPLLRWKRIAVLAAMCIVVVQGII
FQVAYFLHMQTYVLKRPTMLPRSVIFATAFMSFFCLVLALFKDIPDHEGDEKFG
IQSLSVHLGQKRVFWICVSLLEMIYGVAILVGATSSNHWSKMITVSAHAVLAS
VLWYRSKTVDVKSSSATESFYMFLWKLFYAEYLIIPFII*
>CluTAL
MKINNFLGLKDFQKIIIENGKIELDESLLDRVEASFQFLKEFSRNKVIYGVNTGF
GPMAQFKISDEDTHQLQYNLIRSHSSGIGNPLPAQEVKACMLARLNTLSLGNS
GVHQSVVHLLKELINRDITPLIFEHGGVGASGDLVQLAHLALVLIGEGEVFYK
GERTSTKEVFEIEGLEPIQVEIREGLALMNGTSVMSGIGVVNAYKANQLTDISI
KLSCAINEIVQAYDDHLSEALNGTKKHYGQQKVAERMRAHLADSKLIRKRED
HLYTHFEEQEKVFKEKVQEYYSLRCVPQILGPVLDTLEYTETVLENEINSAND
NPIINVEDKHVYHGGNFHGDYISLEMDKLKIVVTKLTMLAERQLNYLLNAKI
NEILPPFVNLGKLGFNFGMQGVQFTATSTTAESQMLSNSMYVHSIPNNNDNQ
DIVSMGTNAAVICRKVIENAFEVLAIEAITIIQAVEYLGFQDKVSSSTKELYDEI
RKIIPAFSNDMVMYPYLEEVKKYLKAM*
>RgPAL
MAPSVDSIATSVANSLSNGLAGDVRKKTSGAGSLLPTTETTQIDIVERILADAG
ATDQIKLDGYTLTLGDVVGAARHGRTVKVADSPQIREKIDASVEFLRTQLDNS
VYGVTTGFGGSADTRTEDAISLQKALLEHQLCGVLPTSMDGFALGRGLENSL
PLEVVRGAMTIRVNSLTRGHSAVRIVVLEALTNFLNHGITPIVPLRGTISASGDL
SPLSYIAASITGHPDSKVHVDGKIMSAQEAIALKGLQPVVLGPKEGLGLVNGT
AVSASMATLALTDAHVLSLLAQANTALTVEAMVGHAGSFHPFLHDVTRPHPT
QIEVARNIRTLLEGSKYAVHHETEVKVKDDEGILRQDRYPLRCSPQWLGPLVS
DMIHAHSVLSLEAGQSTTDNPLIDLENKMTHHGGAFMASSVGNTMEKTRLA
VALMGKVSFTQLTEMLNAGMNRALPSCLAAEDPSLSYHCKGLDIAAAAYTSE
LGHLANPVSTHVQPAEMGNQAINSLALISARRTAEANDVLSLLLATHLYCVLQ
AVDLRAMEFEHTKEFEPMVTALLKQHLGALATADVEDKVRKSIYKRLQQNNS
YDLEQRWHDTFSVATGAVVEALAGNEVSLASLNAWKVACAEKAIALTRTVRD
SFWAAPSSSSPALKYLSPRTRILYSFIREDVGVKARRGDVYLGKQEVTIGTNVS

RIYEAIKDGRIAPVLVKMMA*
>YlCYB5
MIINGKVYDISSFVDEHPGGEEVLLDAGGTEATNAFDDVGHSEDAYGILNDLY
VGEVDPSEDVIRKTHTVKTSYEDGESVGDDHGSSSMIFLIVAAAVAAAAFFYL
QGQK*
>ATR2
MSSSSSSSTSMIDLMAAIIKGEPVIVSDPANASAYESVAAELSSMLIENRQFAMI
VTTSIAVLIGCIVMLVWRRSGSGNSKRVEPLKPLVIKPREEEIDDGRKKVTIFFG
TQTGTAEGFAKALGEEAKARYEKTRFKIVDLDDYAADDDEYEEKLKKEDVAF
FFLATYGDGEPTDNAARFYKWFTEGNDRGEWLKNLKYGVFGLGNRQYEHF
NKVAKVVDDILVEQGAQRLVQVGLGDDDQCIEDDFTAWREALWPELDTILRE
EGDTAVATPYTAAVLEYRVSIHDSEDAKFNDINMANGNGYTVFDAQHPYKAN
VAVKRELHTPESDRSCIHLEFDIAGSGLTYETGDHVGVLCDNLSETVDEALRLL
DMSPDTYFSLHAEKEDGTPISSSLPPPFPPCNLRTALTRYACLLSSPKKSALVAL
AAHASDPTEAERLKHLASPAGKVDEYSKWVVESQRSLLEVMAEFPSAKPPLG
VFFAGVAPRLQPRFYSISSSPKIAETRIHVTCALVYEKMPTGRIHKGVCSTWMK
NAVPYEKSENCSSAPIFVRQSNFKLPSDSKVPIIMIGPGTGLAPFRGFLQERLAL
VESGVELGPSVLFFGCRNRRMDFIYEEELQRFVESGALAELSVAFSREGPTKE
YVQHKMMDKASDIWNMISQGAYLYVCGDAKGMARDVHRSLHTIAQEQGSM
DSTKAEGFVKNLQTSGRYLRDVW*
>EbC4H
MDLLLLEKTLIALFATIIITIIISKLRGKKFKLPPGPTAVPIFGNWLQVGDDLNHR
NLTDLAKKFGQIFLLRMGQRNLVVVSSPDLAKEVLHTQGVEFGSRTRNVVFDI
FTGKGQDMVFTVYGEHWRKMRRIMTVPFFTNKVVQQYRKGWEDEAAAVVE
DVKKNPKSATEGVVIRKRLQLMMYNNMFRIMFDRRFDSEDDPLFLKLKALN
GERSRLAQSFEYNYGDFIPVLRPFLRGYLKLCKEVKDKRLQLFKDYFVDERK
KIGSTKKLDNNQLKCAIDHILEAKDKGEINEDNVLYIVENINVAAIETTLWSIE
WGIAELVNHPEIQAKLRHELDTKLGSGVQITEPDVQNLPYLQAVVKETLRLR
MAIPLLVPHMNLHDAKLGGFDIPAESKILVNAWWLANNPDQWKKPEEFRPER
FLEEEAKVEANGNDFRYLPFGVGRRSCPGIILALPILGITLGRLVQNFELLPPPG
QSKIDTSEKGGQFSLHILKHSTIVAKPRSF*
>tHMGR
MTQSVKVVEKHVPIVIEKPSEKEEDTSSEDSIELTVGKQPKPVTETRSLDDLEA
IMKAGKTKLLEDHEVVKLSLEGKLPLYALEKQLGDNTRAVGIRRSIISQQSNTK
TLETSKLPYLHYDYDRVFGACCENVIGYMPLPVGVAGPMNIDGKNYHIPMAT
TEGCLVASTMRGCKAINAGGGVTTVLTQDGMTRGPCVSFPSLKRAGAAKIWL
DSEEGLKSMRKAFNSTSRFARLQSLHSTLAGNLLFIRFRTTTGDAMGMNMISK
GVEHSLAVMVKEYGFPDMDIVSVSGNYCTDKKPAAINWIEGRGKSVVAEATIP
AHIVKSVLKSEVDALVELNISKNLIGSAMAGSVGGFNAHAANLVTAIYLATGQ
DPAQNVESSNCITLMSNVDGNLLISVSMPSIEVGTIGGGTILEPQGAMLEMLG
VRGPHIETPGANAQQLARIIASGVLAAELSLCSALAAGHLVQSHMTHNRSQAP
TPAKQSQADLQRLQNGSNICIRS*
>IDI
MTTSYSDKIKSISVSSVAQQFPEVAPIADVSKASRPSTESSDSSAKLFDGHDEEQ

IKLMDEICVVLDWDDKPIGGASKKCCHLMDNINDGLVHRAFSVFMFNDRGEL
LLQQRAAEKITFANMWTNTCCSHPLAVPSEMGGLDLESRIQGAKNAAVRKLE
HELGIDPKAVPADKFHFLTRIHYAAPSSGPWGEHEIDYILFVRGDPELKVVANE
VRDTVWVSQQGLKDMMADPKLVFTPWFRLICEQALFPWWDQLDNLPAGDD
EIRRWIK*
>ARO4$^{K221L}$
MSRSSSPNASSAEDVRILGYDPLLAPALLQTEVASTKNARETVSKGRKDSIDVI
TGKSDKLLCIVGPCSLHDPKAAMEYAQRLKELSDKLSGELVIVMRAYLEKPRT
TVGWKGLINDPDMDESFNINKGLRLSRKVFCDLTDLGLPIASEMLDTISPQFL
ADLLSLGAIGARTTESQLHRELASGLSFPVGFKNGTDGTLGVAVDAVQAASHP
HHFMGVTLQGVAAITTTKGNENCFIILRGGKKGTNYDAESVAECKKATESML
MVDCSHGNSNKDYRNQPKVSKAVAEQVAAGEKKIIGVMIESNIHEGNQKVPK
EGPSALKYGVSITDACVSWETTVDMLTELANAVKERRNKN*
>ARO7$^{G139S}$
MDFTKADTVLDLANIRDSLVRMEDTIVFNLIERAQFCRSEFVYKAGNSDIPGF
KGSYLDWFLQESEKVHAKLRRYAAPDEQAFFPDDLPEAILPPIDYAPILAPYSK
EVSVNDEIKKIYTDDIVPLVCAGTGDQPENYSSVMVCDIETLQALSRRIHFGKF
VAESKFLSETERFTELIKNKDIAGIEAAITNSKVEETILARLGEKALAYGTDPTL
RWSQRTQGKVDSEVVKRIYKEWVIPLTKKVEVDYLLRRLE*
>ARO1
MFAEGQIQKVPILGKESIHIGYKMQDHIVSEIVANIKSSTYILVTDTNIEDLGYV
ESLKTKFEAAFAKDGIKSRLLTYTVAPGETSKSRATKAAIEDWMLSKGCTRDT
VILAVGGGVIGDMIGYVAATFMRGVRFVQIPTTLLAMVDSSIGGKTAIDTPLGK
NLVGAFWQPVNIFIDTSFLETLPVREFINGMAEVIKTAAFYDAEEFTRLESASEI
FLSTIKKRDAKDPRRVDLSPITDTIGRIVLGSARIKAAVVSADEREGGLRNLLN
FGHSIGHAYEAILTPYILHGECVAIGMVKEAELSRYLGILSPVAVARLAKCIKAY
ELPVSLDDATVKARSHGKKCPVDDLLRIMGVDKKNDGSTKKIVILSAIGKTHE
QKASSVADKDIRFVLSEEVIVGEAPVGDKKSYTVTPPGSKSISNRAFVLTALGK
GPCKLRNLLHSDDTQHMLEAIELLGGASFEWEADGETLLVTGNGGKLTAPAQ
ELYLGNAGTASRFLTTAATLVQKGDKDHVILTGNKRMQERPIGPLVDALRSNG
ADIAFQNAEGSLPLKIEAGVGLKGGLIEVAATVSSQYVSSLLMCAPYAQTPVT
LSLVGGKPISQFYIDMTIAMMADFGVVVTKDETKEHTYHIPQGVYTNPEEYV
VESDASSATYPLAYAAMTGHTVTVPNIGSKSLQGDARFAIDVLKAMGCTVEQ
TATSTTVTGVPNLKAIAVDMEPMTDAFLTACVVAAVSEGTTVITGIANQRVKE
CNRIEAMRVQLAKYGVVCRELEDGIEVDGISRSDLKTPVSVHSYDDHRVAMS
FSLLSSIMAAPVAIEERRCVEKTWPGWWDVLSGVFNVPLEGVTLAKTVSKAE
SGLSKPSIFIVGMRGAGKTHLGAQAANHLGYEFIDLDQLLEKDLDTTIPQLIAD
KGWDHFRAEELRLLKQCLNDKSEGYVISCGGGVVETPAARDALQTFKGVGGI
VLHVHRPVSRILEYLNKDQSRPAFVDDLEAVWQRRKELYRSVSSNVFFAPHC
DSAEATAKVQQMLGAFLDRVTGKSEFVIPHKDQFTSFLSLTFPDVSIAATMLPS
LSEGCSALELRVDLLNENDEAIPSEEYVLSQLAILRQNVDLPILYTVRTKAQGG
RFPDDKPVELANLVNLGLKTAVELLDVELTYPAELVSSVGASRGYTKLLGSHH
DFPGALNWSSLEWENMYARAEAVPVDVVKLVGMAKSFSDNFALENFREAHT
SSPLLAINMGSHGQLSRVTNTLLTPVTHADLPVAAAPGQLSVEEINQTRSTIGM

FNKNLSFFIVGTPIGHSKSPILHNTMFKKLGLPYEYSRFKTDDAAAVNAKARA
LLAQGNLGGISVTIPLKQDIIPFLDEVSPLAQQIGAVNTIIPGPNGTLKGDNTDIL
GLVNALTRFGANSLDKKTALIVGAGGTSLAAVHGLRSLGFAKILIANRTLSKAE
AIADKFDNVEAVTLDSFVANKYTPSVIVSCVPATTFSMLDESNKLVSAALAASP
KGLVLEAAYSAEATPLLKQVMDVEGWEFISGLYMLTEQGFEQFRLWTGIPAPK
EVGEKAVLGN*
>ARO2
MSTFGTLFKVTTYGESHCKSVGCIVDGVPPGMDLDESDIQPQLTRRRPGQSAL
TTPRNERDAVAIQSGTEYGKTLGTPIAMLVQNKDQRPHDYSEMDDYPRPSHA
DYTYQEKYGIKASSGGGRSSARETIGRVAAGAIADKYLAAVNDIEIVAFVSQV
GDVSMDRSPRNEQWISTLEGVTREGIDSTGPMRCPDLALGEKMVKIVEEHRD
SHDSVGGVVTCVIRNCPVGLGEPCFDKLEATLAHAMMSIPATKGFEFGSGFAG
AAMSGSKHNDMFYKDVASGRFRTRTNYSGGVQGGISNGENIYFNIAFKPPATI
SQEQATATYAGKDGVLAAKGRHDPNVVPRAVPIVEAMAALVIADAHLIQESR
KGAKSFF*
>ARO3$^{K225L}$
MPAMHNASNAQGDRNRTEDWRIRGYNPLTAPDLLQHEIPLTKQSKATILKGR
QDACDILDGKDDRIIVVVGPCSIHDPKAAMEYAERLKQISDKLSGELLIVMRA
YLEKPRTTVGWKGLINDPDMDGSFKINKGLRVARDLFVKLTELNPIASELLDTI
SPQFLADLFSVGAIGARTTESQLHRELASGLSFPVGFKNGTDGGIKVALDAIQA
AAHPHHFLSVTLPGVVAIVGTDGNEDCFLILRGGSKGPNYDAEHVAEVKKQV
GETKGPRIMVDCSHGNSSKNHKNQPLVASNVAQQIAAGEKSICGLMIESNIHE
GRQDICDNKEDMKYGVSVTDACINWEDTEKVLEELAQAVKTRRG*
>AroG*
MNYQNDDLRIKEIKELLPPVALLEKFPATENAANTVAHARKAIHKILKGNDDR
LLVVIGPCSIHDPVAAKEYATRLLALREELKDELEIVMRVYFEKPRTTVGWKG
LINDPHMDNSFQINDGLRIARKLLLDINDSGLPAAGEFLNMITLQYLADLMSW
GAIGARTTESQVHRELASGLSCPVGFKNGTDGTIKVAIDAINAAGAPHCFLSVT
KWGHSAIVNTSGNGDCHIILRGGKEPNYSAKHVAEVKEGLNKAGLPAQVMID
FSHANSSKQFKKQMDVCADVCQQIAGGEKAIIGVMVESHLVEGNQSLESGEP
LAYGKSITDACIGWEDTDALLRQLANAVKARRG*
>TyrA*
MVAELTALRDQIDEVDKALLNLLAKRLELVAEVGEVKSRFGLPIYVPEREASIL
ASRRAEAEALGVPPDLIEDVLRRVMRESYSSENDKGFKTLCPSLRPVVIVGGG
GQMGRLFEKMLTLSGYQVRILEQHDWDRAADIVADAGMVIVSVPIHVTEQVI
GKLPPLPKDCILVDLASVKNGPLQAMLVAHDGPVLGLHPMFGPDSGSLAKQV
VVWCDGRKPEAYQWFLEQIQVWGARLHRISAVEHDQNMAFIQALRHFATFAY
GLHLAEENVQLEQLLALSSPIYRLELAMVGRLFAQDPQLYADIIMSSERNLALI
KRYYKRFGEAIELLEQGDKQAFIDSFRKVEHWFGDYRQRFQSESRVLLRQAN
DNRQ*
>MAF1
MQFIQDHTLDEVASALKFDTPECHVSGNIDLYTTKPASSDKKLYKVLDKQLSY
HQDALDSFTSISTLGTSFSPPQVLTFDNYYQYGKSLDSSVSKMSNRSPKGVSK
VPKQRTYSSSRAGVSVGSLSPLLSSPFGDMSETASRKVFAYLIAILNASDPNHD

FSVLQPDDFQREPSASAVISSFNNVLFGLGMPIPPRMWDVLDKSIEMHNCTIFS
FTPDATILADEPGALWALSWFFFNKRMKRVACISLNARRHSLSPTLGPKIMRSD
VINGDDEEDYDLTYSSDTDMYDEIVGELDLE*
>AgrGPPS2
MAYSAMATMGYNGMAASCHTLHPTSPLKPFHGASTSLEAFNGEHMGLLRGY
SKRKLSSYKNPASRSSNATVAQLLNPPQKGKKAVEFDFNKYMDSKAMTVNEA
LNKAIPLRYPQKIYESMRYSLLAGGKRVRPVLCIAACELVGGTEELAIPTACAI
EMIHTMSLMHDDLPCIDNDDLRRGKPTNHKIFGEDTAVTAGNALHSYAFEHIA
VSTSKTVGADRILRMVSELGRATGSEGVMGGQMVDIASEGDPSIDLQTLEWI
HIHKTAMLLECSVVCGAIIGGASEIVIERARRYARCVGLLFQVVDDILDVTKSS
DELGKTAGKDLISDKATYPKLMGLEKAKEFSDELLNRAKGELSCFDPVKAAP
LLGLADYVAFRQN*
>GgaFPS$^{N144W}$
MHKFTGVNAKFQQPALRNLSPVVVEREREEFVGFFPQIVRDLTEDGIGHPEVG
DAVARLKEVLQYNAPGGKCNRGLTVVAAYRELSGPGQKDAESLRCALAVGW
CIELFQAFFLVADDIMDQSLTRRGQLCWYKKEGVGLDAIWDSFLLESSVYRVL
KKYCRQRPYYVHLLELFLQTAYQTELGQMLDLITAPVSKVDLSHFSEERYKAI
VKYKTAFYSFYLPVAAAMYMVGIDSKEEHENAKAILLEMGEYFQIQDDYLDC
FGDPALTGKVGTDIQDNKCSWLVVQCLQRVTPEQRQLLEDNYGRKEPEKVAK
VKELYEAVGMRAAFQQYEESSYRRLQELIEKHSNRLPKEIFLGLAQKIYKRQK
*
>ERG10
MRLTLPRLNAAYIVGAARTPVGKFNGALKSVSAIDLGITAAKAAVQRSKVPAD
QIDEFLFGQVLTANSGQAPARQVVIKGGFPESVEATTINKVCSSGLKTVALAAQ
AIKAGDRNVIVAGGMESMSNTPYYSGRGLVFGNQKLEDSIVKDGLWDPYNNI
HMGNCCENTNKRDGITREQQDEYAIESYRRANESIKNGAFKDEIVPVEIKTRK
GTVTVSEDEEPKGANAEKLKGLKPVFDKQGSVTAGNASPINDGASAVVVASG
TKAKELGTPVLAKIVSYADAATAPIDFTIAPSLAIPAALKKAGLTKDDIALWEIN
EAFSGVALANLMRLGIDKSKVNVKGGAVALGHPIGASGNRIFVTLVNALKEGE
YGVAAICNGGGASTAIVIKKVSSVE*
>ERG13
MSQPQNVGIKALEIYVPSRIVNQAELEKHDGVAAGKYTIGLGQTNMAFVDDR
EDIYSFALTAVSRLLKNNNIDPASIGRIEVGTETLLDKSKSVKSVLMQLFGENSN
IEGVDNVNACYGGTNALFNAINWVEGRSWDGRNAIVVAGDIALYAKGAARP
TGGAGCVAMLIGPDAPLVLDNVHGSYFEHAYDFYKPDLTSEYPYVDGHYSLT
CYTKALDKAYAAYNARAEKVGLFKDSDKKGADRFDYSAFHVPTCKLVTKSY
ARLLYNDYLNDKSLYEGQVPEEVAAVSYDASLTDKTVEKTFLGIAKAQSAER
MAPSLQGPTNTGNMYTASVYASLISLLTFVPAEQLQGKRISLFSYGSGLASTLF
SLTVKGDISPIVKACDFKAKLDDRSTETPVDYEAATDLREKAHLKKNFEPQGD
IKHIKSGVYYLTNIDDMFRRKYEIKQ*
>ERG19
MIHQASTTAPVNIATLKYWGKRDPALNLPTNNSISVTLSQDDLRTLTTASCSPD
FTQDELWLNGKQEDVSGKRLVACFRELRALRHKMEDSDSSLPKLADQKLKIV
SENNFPTAAGLASSAAGFAALIRAVANLYELQETPEQLSIVARQGSGSACRSLY

GGYVAWEMGTESDGSDSRAVQIATADHWPEMRAAILVVSADKKDTSSTTGM
QVTVHTSPLFKERVTTVVPERFAQMKKSILDRDFPTFAELTMRDSNQFHATCL
DSYPPIFYLNDVSRASIRVVEAINKAAGATIAAYTFDAGPNCVIYYEDKNEELV
LGALKAILGRVEGWEKHQSVDAKKIDVDERWESELANGIQRVILTKVGGDPV
KTAESLINEDGSLKNSK*
>ERG8
MTTYSAPGKALLCGGYLVIDPAYSAYVVGLSARIYATVSASEASTTSVHVVSP
QFDKGEWTYNYTNGQLTAIGHNPFAHAAVNTVLHYVPPRNLHINISIKSDNAY
HSQIDSTQRGQFAYHKKAIHEVPKTGLGSSAALTTVLVAALLKSYGIDPLHNT
HLVHNLSQVAHCSAQKKIGSGFDVASAVCGSLVYRRFPAESVNMVIAAEGTSE
YGALLRTTVNQKWKVTLEPSFLPPGISLLMGDVQGGSETPGMVAKVMAWRK
AKPREAEMVWRDLNAANMLMVKLFNDLRKLSLTNNEAYEQLLAEAAPLNA
LKMIMLQNPLGELARCIITIRKHLKKMTRETGAAIEPDEQSALLNKCNTYSGVI
GGVVPGAGGYDAISLLVISSTVNNVKRESQGVQWMELKEENEGLRLEKGFK*
>ERG12
MDYIISAPGKVILFGEHAAVFGKPAIAAAIDLRTYLLVETTTSDTPTVTLEFPDI
HLNFKVQVDKLASLTAQTKADHLNWSTPKTLDKHIFDSLSSLALLEEPGLTKV
QQAAVVSFLYLYIHLCPPSVCEDSSNWVVRSTLPIGAGLGSSASICVCLAAGLL
VLNGQLSIDQARDFKSLTEKQLSLVDDWSFVGEMCIHGNPSGIDNAVATQGGA
LLFQRPNNRVPLVDIPEMKLLLTNTKHPRSTADLVGGVGVLTKEFGSIMDPIMT
SVGEISNQAMEIISRGKKMVDQSNLEIEQGILPQPTSEDACNVMEDGATLQKL
RDIGSEMQHLVRINHGLLIAMGVSHPKLEIIRTASIVHNLGETKLTGAGGGGCA
ITLVTSKDKTATQLEENVIAFTEEMATHGFEVHETTIGARGVGMCIDHPSLKTV
EAFKKVERADLKNIGPWTH*
